# The role of inhibitory neurons in deviance sound detection in regular and random statistical contexts

**DOI:** 10.1101/2025.04.20.649735

**Authors:** Xiaomao Ding, Nathan W. Vogler, Melanie Tobin, Linda Garami, Alexandria M.H. Lesicko, Katherine C. Wood, Maria N. Geffen

## Abstract

Detecting statistical regularities in sound and responding to violations of these patterns, termed deviance detection, is a core function of the auditory system. In the human brain, studies have shown that deviance responses are enhanced in regular compared to random auditory contexts, but the underlying neuronal circuit mechanisms remain unclear. Here, we examined how inhibitory neurons contribute to context-dependent deviance responses in mouse auditory cortex (AC). Using two-photon calcium imaging in AC of awake head-fixed male and female mice, we recorded neuronal activity during presentation of spectro-temporally rich moving ripple sounds, with deviant ripples embedded in either regular or random ripple sequences. AC neurons exhibited stronger responses to deviant sounds in regular contexts compared to random ones. To identify circuit mechanisms, we selectively inactivated parvalbumin (PV), somatostatin (SST), or vasoactive intestinal polypeptide (VIP) inhibitory neurons during the deviant stimulus presentation. Inactivation of PV and SST neurons broadly increased deviance responses in both contexts. In contrast, VIP inactivation selectively reduced responses to deviant stimuli in the regular, but not random, context, decreasing the context-dependent deviance signal enhancement. At the population level, inactivating all three neuronal subtypes increased detectability of the deviant stimulus, but the effects were context-dependent only for VIP inactivation. These findings reveal a distinct role for VIP neurons in modulating deviance signals based on context regularity, identifying a specific inhibitory neuron type that is critical for context-sensitive auditory processing and predictive coding.

**SIGNIFICANCE STATEMENT:** Understanding how the brain detects and responds to predictable and unexpected sounds is critical for communication and navigation in complex acoustic environments. Although prior studies have shown that detection of novel sounds is enhanced by regularity in background sound patterns, the underlying circuit mechanisms have remained unknown. Here, we find that neurons in the auditory cortex exhibit stronger deviance responses when the deviant sounds are embedded in regular, predictable sound sequences as compared to random sound sequences. Importantly, we identify a distinct role for specific inhibitory neurons in modulating this context-dependent enhancement. These findings reveal a new insight into how cortical circuits implement predictive coding strategies to optimize sensory processing.

## INTRODUCTION

Successfully navigating everyday sound environments requires the auditory system to detect and interpret unexpected sounds within predictable patterns (Barascud et al., 2016; Sohoglu and Chait, 2016; Parras et al., 2017; Southwell and Chait, 2018). Natural sounds are characterized not only by their spectral content but also by higher-order spectro-temporal regularities: statistical relations that allow listeners to form predictions about upcoming auditory events. Detecting violations of these predictions—referred to as *deviance detection*—is a core auditory computation, supporting a wide range of behaviors, from identifying a familiar voice in a noisy crowd to detecting an approaching vehicle amid ambient noise (Arnal and Giraud, 2012). The human brain encodes both predictions and prediction errors, with signals such as the mismatch negativity (MMN) reflecting changes in neuronal activity in response to unexpected sounds (Naatanen et al., 1978, 2004, 2007). Deviance signals have been observed across multiple stages of the auditory pathway, including the inferior colliculus (IC) (Lesicko et al., 2022), medial geniculate body (MGB) (Antunes et al., 2010; Malmierca et al., 2015), and auditory cortex (AC) (Nelken, 2004, 2014; Hershenhoren et al., 2014; Natan et al., 2015, 2017; Rubin et al., 2016; Parras et al., 2017). However, the transformations of prediction and deviance signals along this pathway, and the neural mechanisms that shape their context sensitivity, remain poorly understood.

In humans, deviance responses are enhanced when deviant sounds are embedded within regular contexts (e.g., a predictable sequence ABCD, ABCD…) as compared to random ones (e.g. shuffled sequence: BADC, ADCB…) (Sohoglu and Chait, 2016; Southwell and Chait, 2018). Stronger responses to a deviant sound following a regular pattern suggest that the brain modulates prediction strength based on context. Within AC, nearly all neurons adapt to frequently presented sounds and an enhanced response to rare, deviant stimuli (Harms et al., 2014; Parras et al., 2017) — a phenomenon known as stimulus-specific adaptation (SSA) (Ulanovsky et al., 2003; Szymanski et al., 2009; Farley et al., 2010; Natan et al., 2015). AC neurons are also sensitive to patterns with structure across multiple time scales (Ulanovsky et al., 2004; Yaron et al., 2012; Mehra et al., 2022). Here, we asked whether individual AC neurons exhibit enhanced deviance responses in regular as compared to random contexts, and whether intra-cortical inhibitory neurons support this transformation.

Inhibitory neurons in AC play distinct roles in shaping auditory cortical responses to sound patterns. The two most common subtypes, parvalbumin (PV)- and somatostatin (SST)-expressing neurons (Lee et al., 2010; Tremblay et al., 2016; Wood et al., 2017), differentially contribute to SSA and other forms of auditory adaptation (Levy and Reyes, 2012; Chen et al., 2015; Natan et al., 2015, 2017; Phillips et al., 2017), while another prominent subtype, vasoactive intestinal polypeptide (VIP)-expressing neurons, contribute to context- and state-dependent modulation (Lee et al., 2010; Tremblay et al., 2016; Wood et al., 2017). VIP neurons gate top-down influences and can modulate predictive coding processes through a disinhibitory circuit (Pfeffer et al., 2013; Pi et al., 2013; Mesik et al., 2015; Bastos et al., 2023; Ferguson et al., 2023; Najafi et al., 2025). We hypothesized that these distinct inhibitory neuron classes differentially contribute to deviance detection depending on regular or random context.

Using two-photon calcium imaging in awake mice, we recorded neuronal activity of large populations of neurons in AC during presentation of deviant spectro-temporally complex ripple stimuli embedded in either regular or random sound contexts. We optogenetically inactivated PV, SST, or VIP neurons to determine their respective roles in modulating deviance responses. Regular context enhanced deviance responses compared to random context. This enhancement was differentially shaped by distinct inhibitory neuron classes. PV and SST inactivation increased responses to deviant sounds, but the role of context was non-specific, whereas VIP inactivation selectively reduced deviance responses in the regular, but not random context. These results identify VIP neurons as a key component of the circuit that integrates context statistics to shape deviance responses.

## METHODS

### Animals

A total of 14 mice (male/female) were used in this study. The strains used were: Cdh23 x PV-Cre (B6.CAST-Cdh23Ahl+/Kjn, JAX: 002756 x Pvalbtm1(cre)Arbr/J, JAX: 017320; n = 4), Cdh23xSST-Cre (B6.CAST-Cdh23Ahl+/Kjn x Ssttm2.1(cre)Zjh/J, 113 JAX: 013044; n = 4), VIP-IRES-Cre (B6.CAST-Cdh23Ahl+/Kjn x Viptm1(cre)Zjh/J, JAX: 010908; n = 3), VGAT-Cre crossed with transgenic GCaMP8m (Slc32a1tm2(cre)Lowl/J, JAX: 016962 x Igs7tm1(tetO-GCaMP8m,CAG-tTA2)Genie/J, JAX:037718, n = 3), Cdh23 (B6N(Cg)-Cdh23tm2.1Kjn/Kjn, n = 5), aged 12-26 weeks. Cdh23 mice were used because they do not exhibit age-related hearing loss which is common in C57BL/6J mice (Johnson et al., 2017). Mice were housed individually after cranial window implant and were housed on a reversed light-dark cycle. Experimental procedures were performed in accordance with NIH guidelines and approved by the IACUC at the University of Pennsylvania.

### Surgery Procedures

Cranial window implants were implanted following a procedure described previously (Wood et al., 2022; Tobin et al., 2024). Mice were anesthetized with 3% isoflurane which was steadily reduced to 1.5% through the surgery. A 3 mm craniotomy was performed over the left auditory cortex with a 3 mm biopsy punch. A viral 750 nL cocktail (4:1 ratio) of virus encoding GCAMP (7f: AAV1-Syn-jGCaMP7f-WPRE, Addgene 104488 or 8m: pGP-AAV-syn-jGCaMP8m-WPRE, Addgene 162375) and virus encoding Jaws in a Cre-dependent fashion (AAV8-CAG-FLEX-JAWS-kgc-tdTomato, UNC Vector Core or AAV8-CAG-FLEX-JAWS-kgc-tdTomato, Penn Vector Core, Addgene 84446) was injected at 3 locations in the craniotomy, separated 0.3-0.5 mm apart in the anterior-posterior direction. The craniotomy was then sealed with a custom round glass window consisting of a 3 mm window glued to a 4 mm window with Norland Optical Adhesive 68. A custom-made metal headplate was attached to the skull with C&B Metabond dental cement. For transgenic GCaMP8m mice, no virus was injected and only the window implant was performed. Mice were allowed to recover for 3 weeks before imaging started.

### Two-photon imaging

All imaging sessions were performed in a custom-built acoustic isolation booth using previously published methods (Betzel et al., 2019; Wood et al., 2022; Tobin et al., 2024). Mice were habituated to being head fixed for 3 days prior to imaging. Wide-field imaging was performed on each mouse to identify sound-responsive areas of the window before two-photon imaging by presenting 3 tone pips (5kHz, 15kHz and 30 kHz) and mapping the gradients of frequency tuning and regions of reversal (Romero et al., 2020). We imaged in layer 2/3 of auditory cortex of awake head-fixed mice using a two-photon (2P) microscope (Ultima 2Pplus *in vivo* multiphoton microscope, Bruker) with a 940 nm laser (Chameleon Ti-Sapphire) with a 16X Nikon objective with a 0.8 numerical aperture (Thorlabs, N16XLWD-PF) (Figure 1 A, B, D). Recordings were made at 512 x 512 pixels and at 30 frames per second.

**Figure 1.**
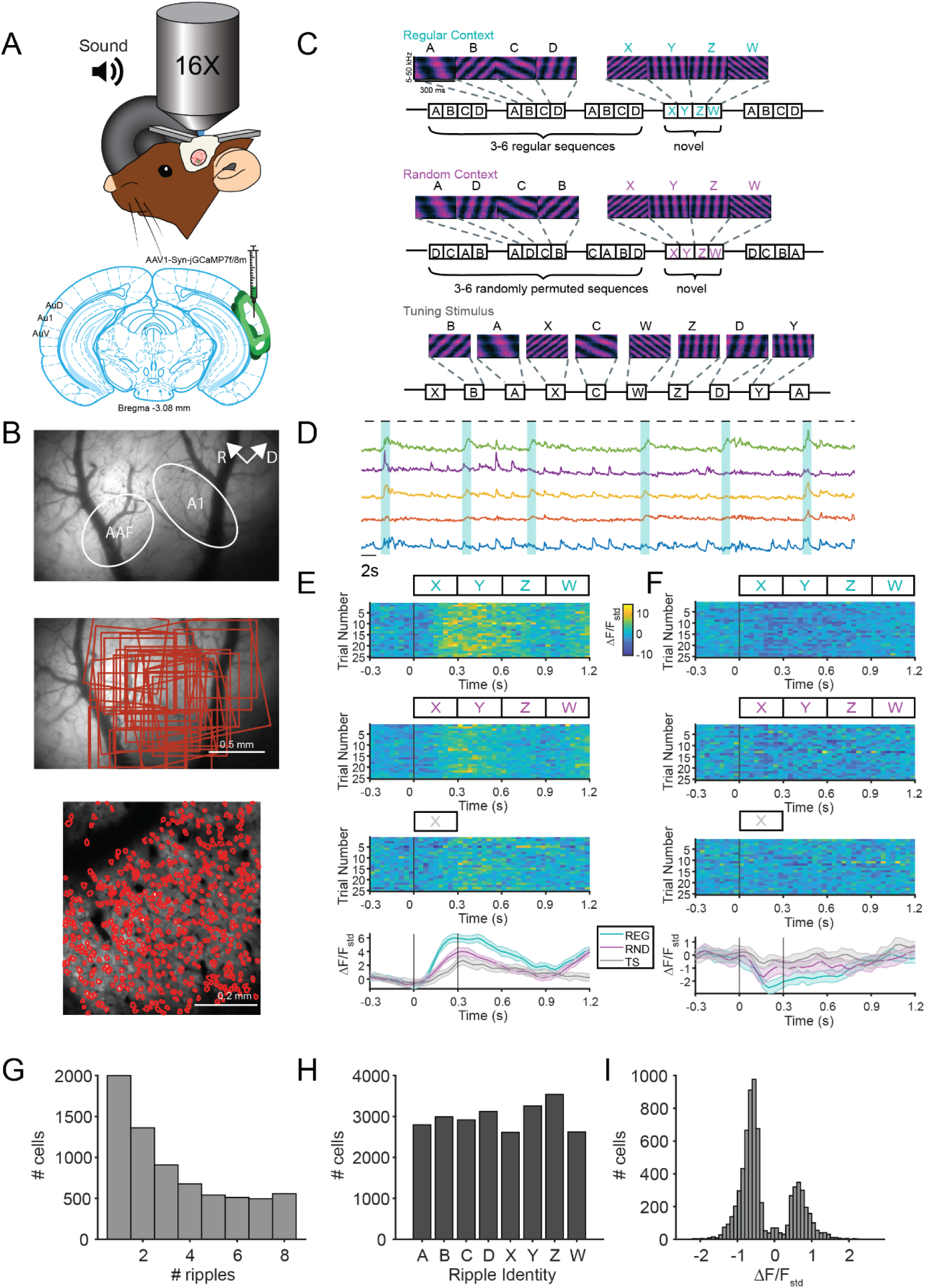
**A.** Schematic of two-photon (2P) calcium imaging setup (top), and viral spread across AC for GCaMP injections (bottom). **B.** Widefield image from an example mouse with estimated areas for A1 and anterior auditory field (AAF) regions (top). Recording sites across different mice mapped onto the example widefield image (middle). Example 2P imaging site with ROIs circled in red (bottom). **C.** Spectrograms of spectro-temporal moving ripples used in the experiment and stimulus schematic. Ripples spanned 5-50 kHz and were 300 ms long in duration. Regular sequences had a fixed temporal pattern (ABCD ABCD…) while random sequences shuffled the background ripple order (DCAB BACD…). The deviant stimulus (novel) sequence was the same for both conditions (XYZW). Each letter in the diagram represents a different spectro-temporal moving ripple. **D.** Example 2P calcium traces from 5 neurons from the same recording session. Black bars on top denote the timing of each ripple sequence. Blue outlines denote the timing of deviant ripple sequences. **E,F.** Responses over the successive trials (top 3 plots) and mean response trace (bottom) to the same deviant ripple in regular (cyan) and random (magenta) contexts as well as the same ripple in the tuning sequence (gray) from example cells with a positive response (**E**) and a negative response (**F**). Shaded error bars represent SEM. **G.** Histogram of number of neurons responsive to only 1 ripple up to all 8 ripples. A significant response was determined with a t-test of the mean response from 0-300 ms post-stimulus a p < 0.05 with a Bonferroni correction. **H.** Histogram of number of neurons responsive to specific ripples. **I.** Histogram of response strengths in response to baseline moving ripples for all sound responsive neurons.

**Table 1.**
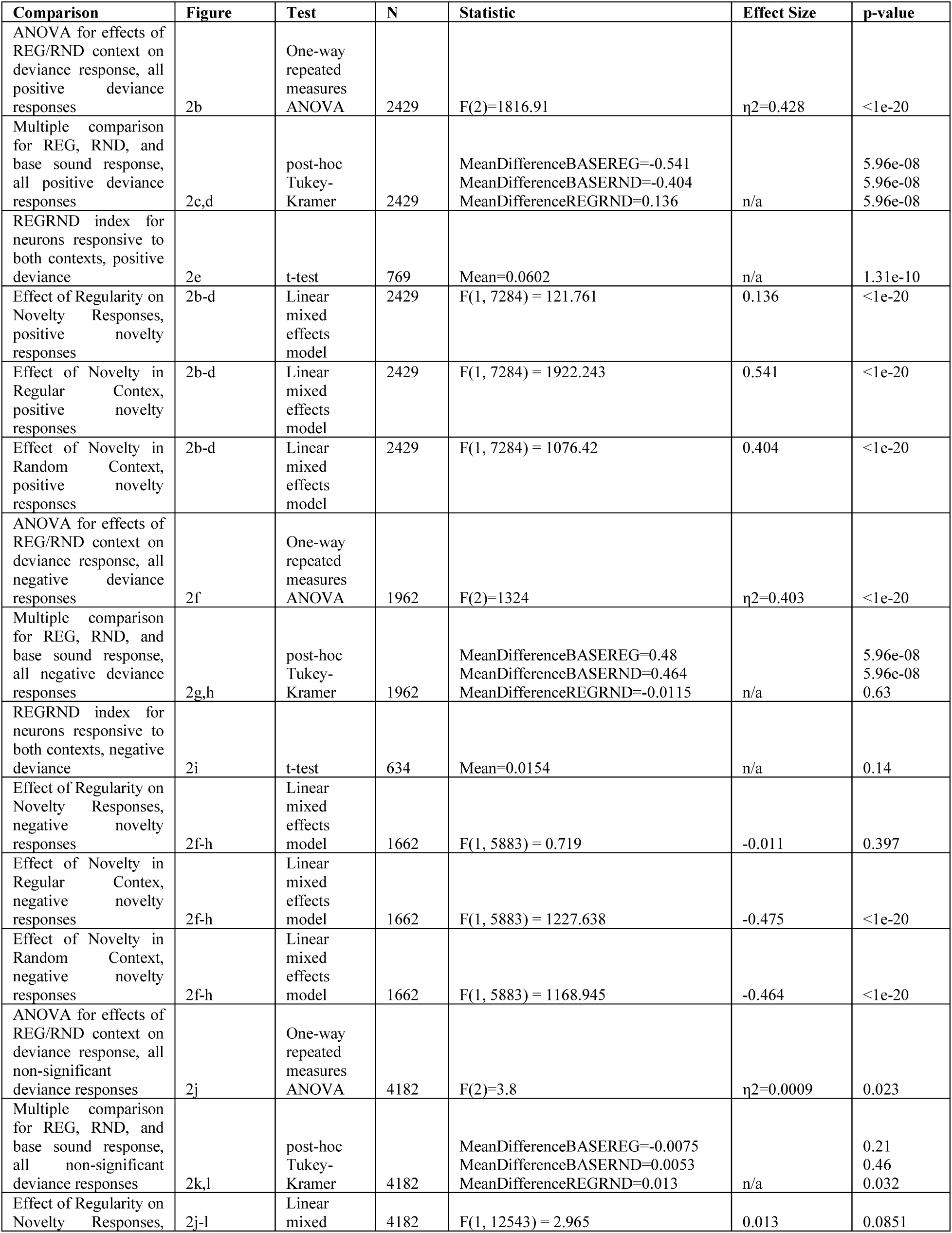
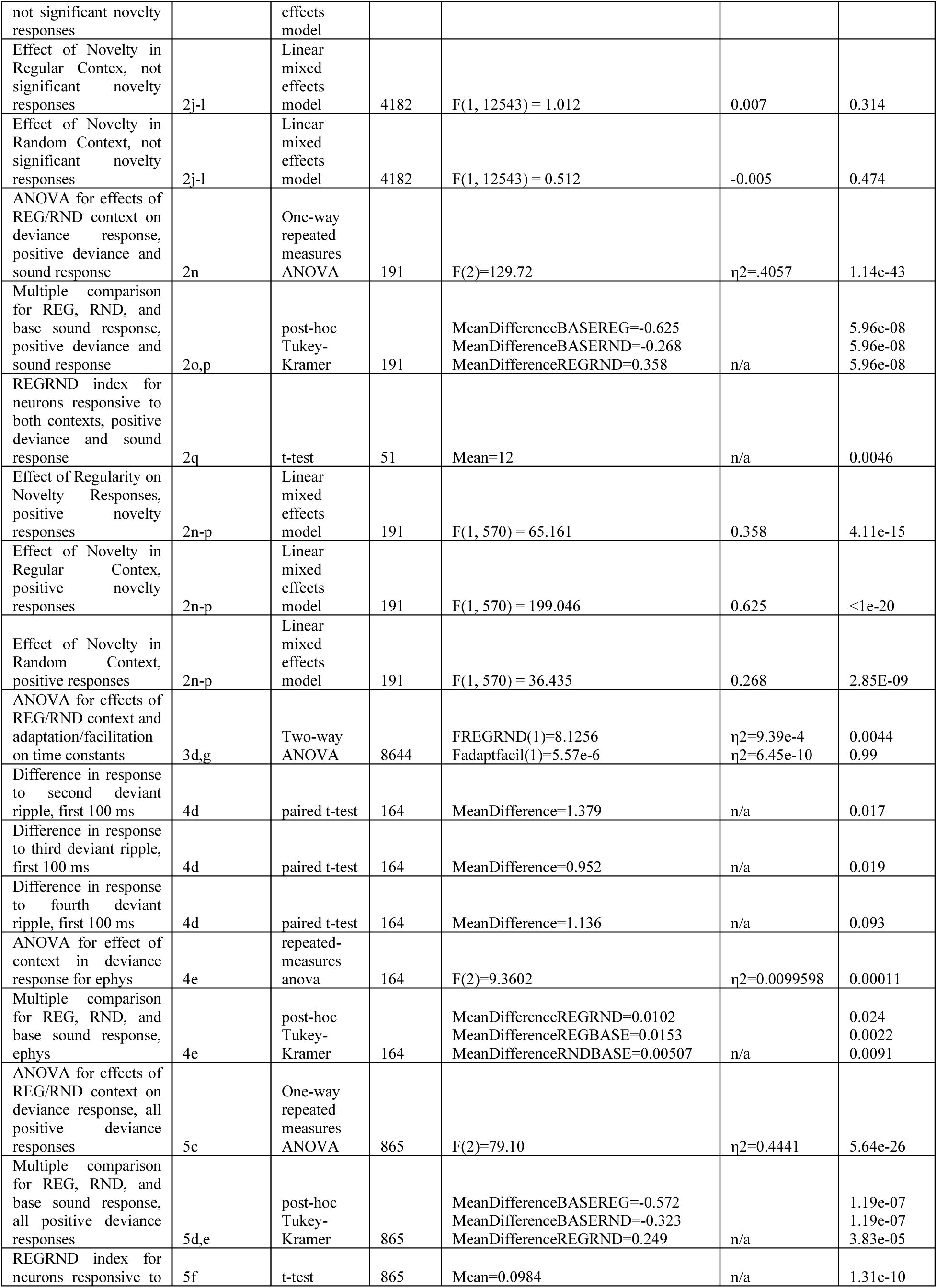
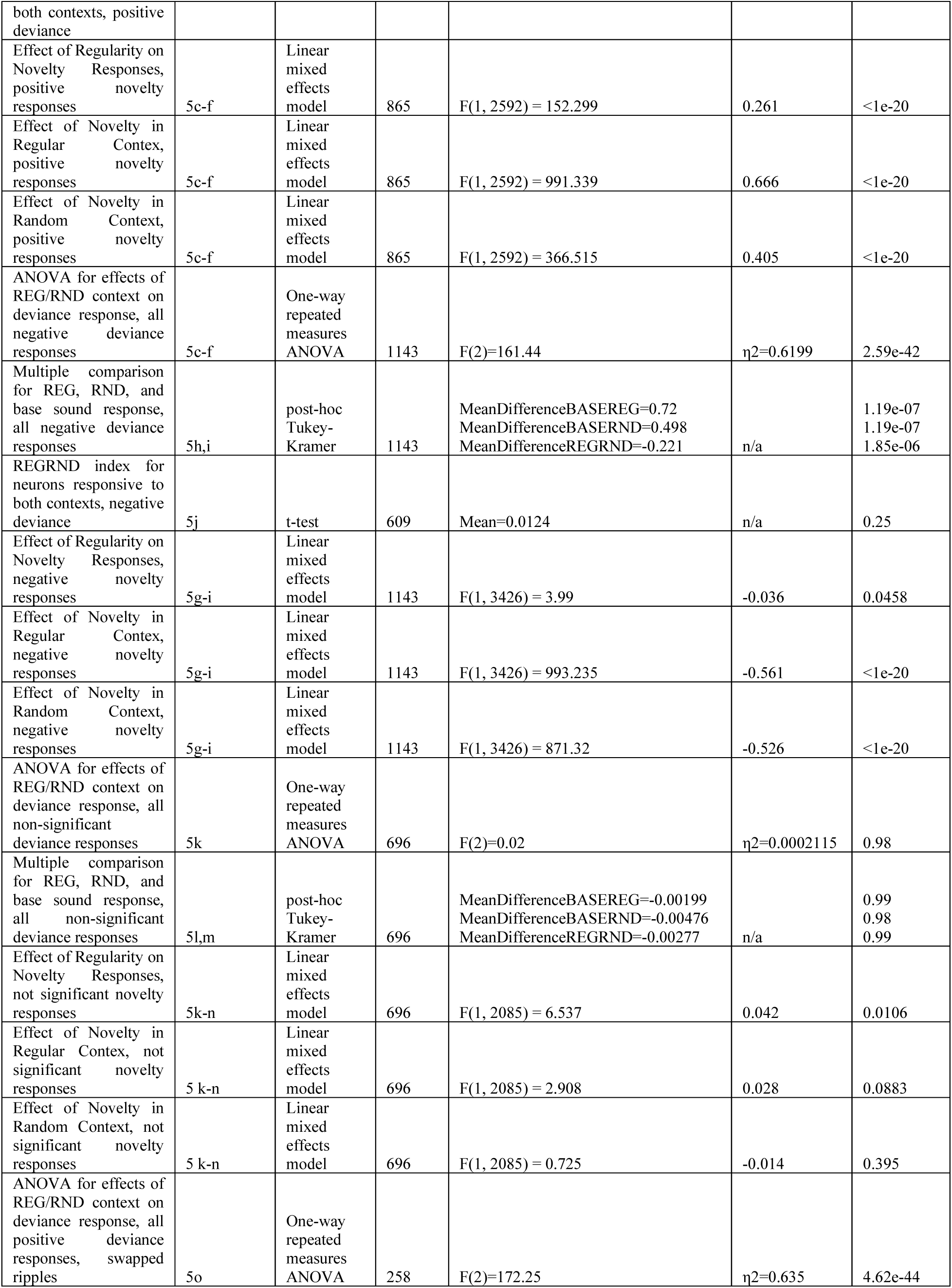
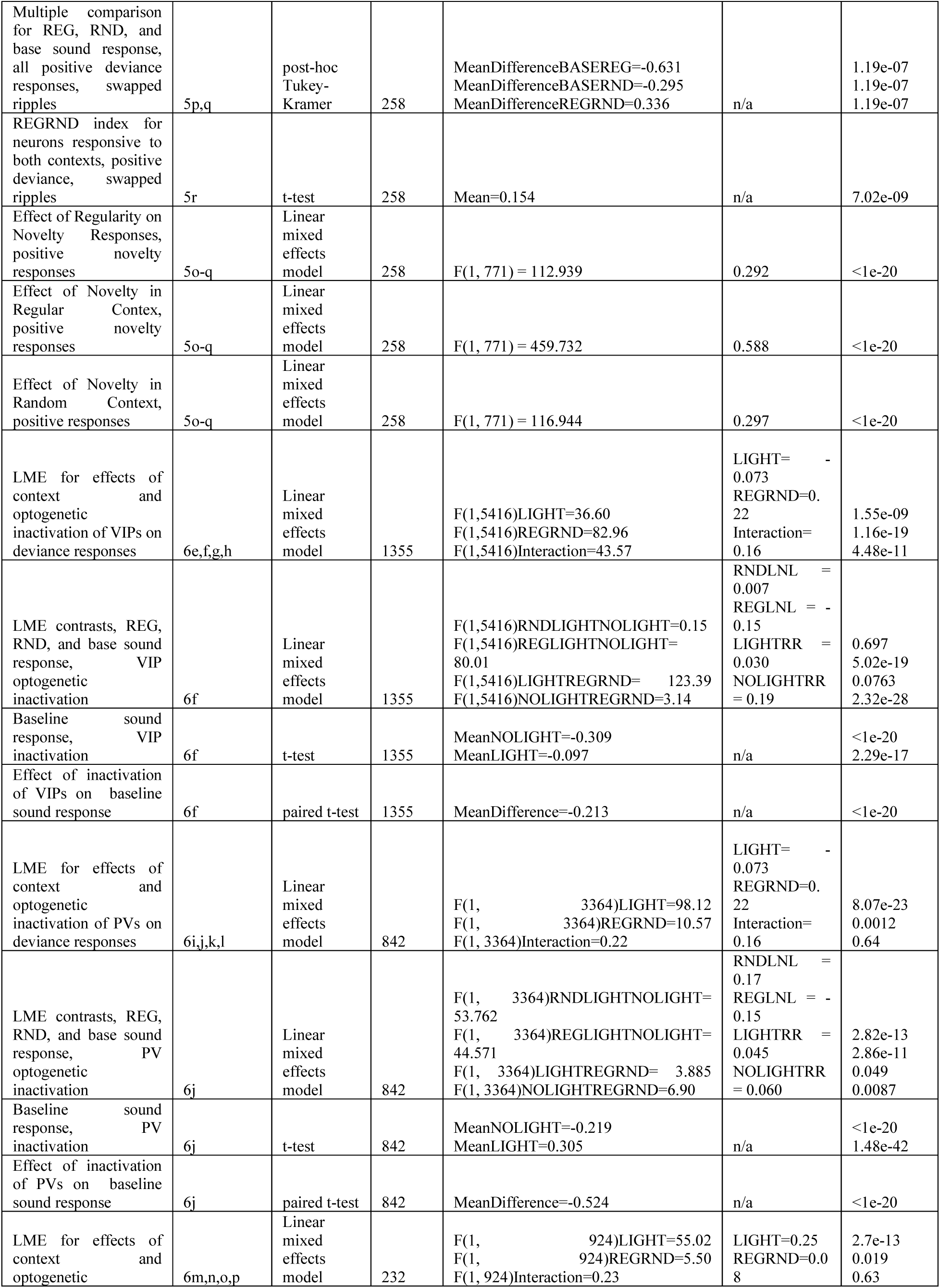
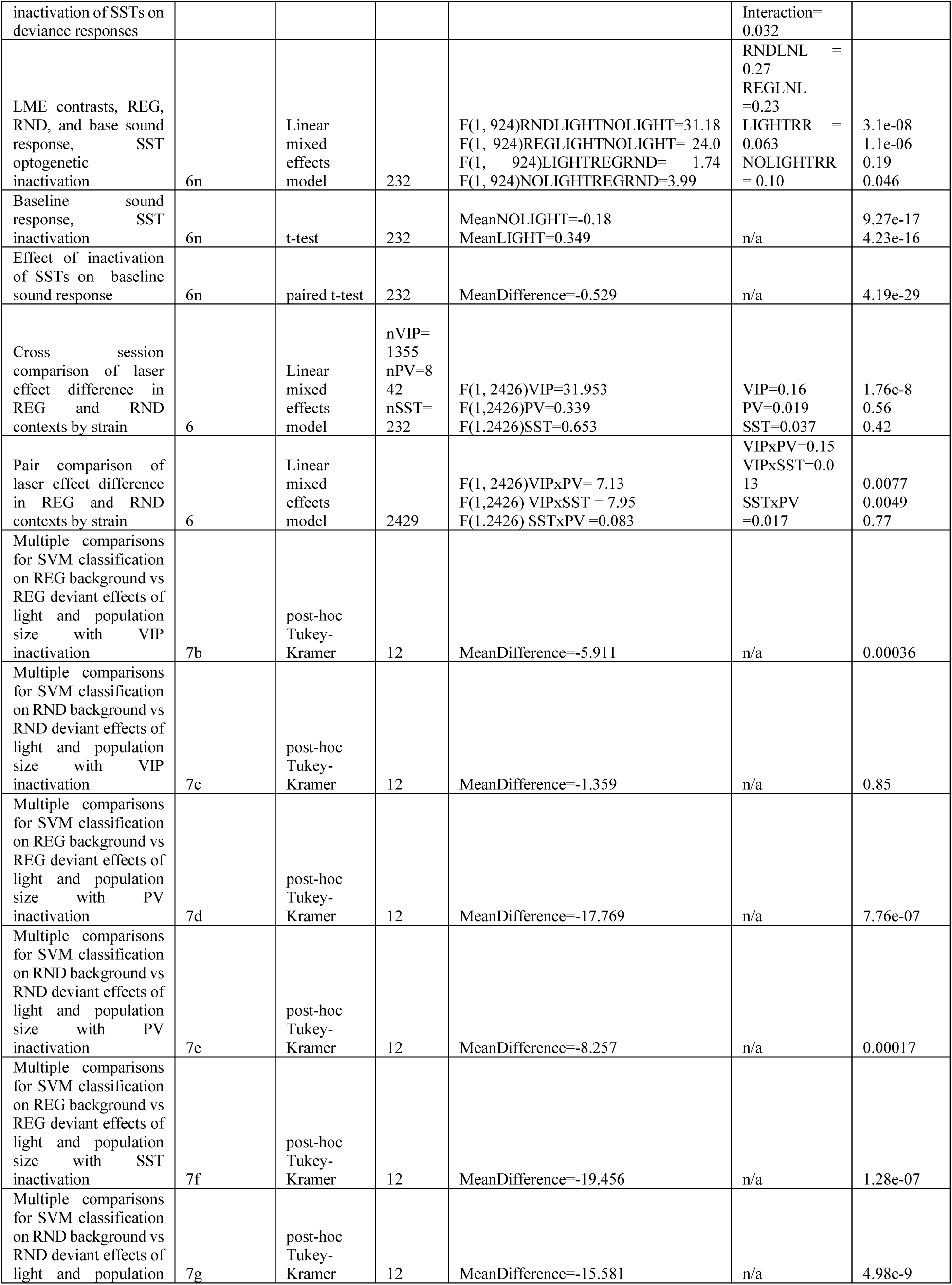
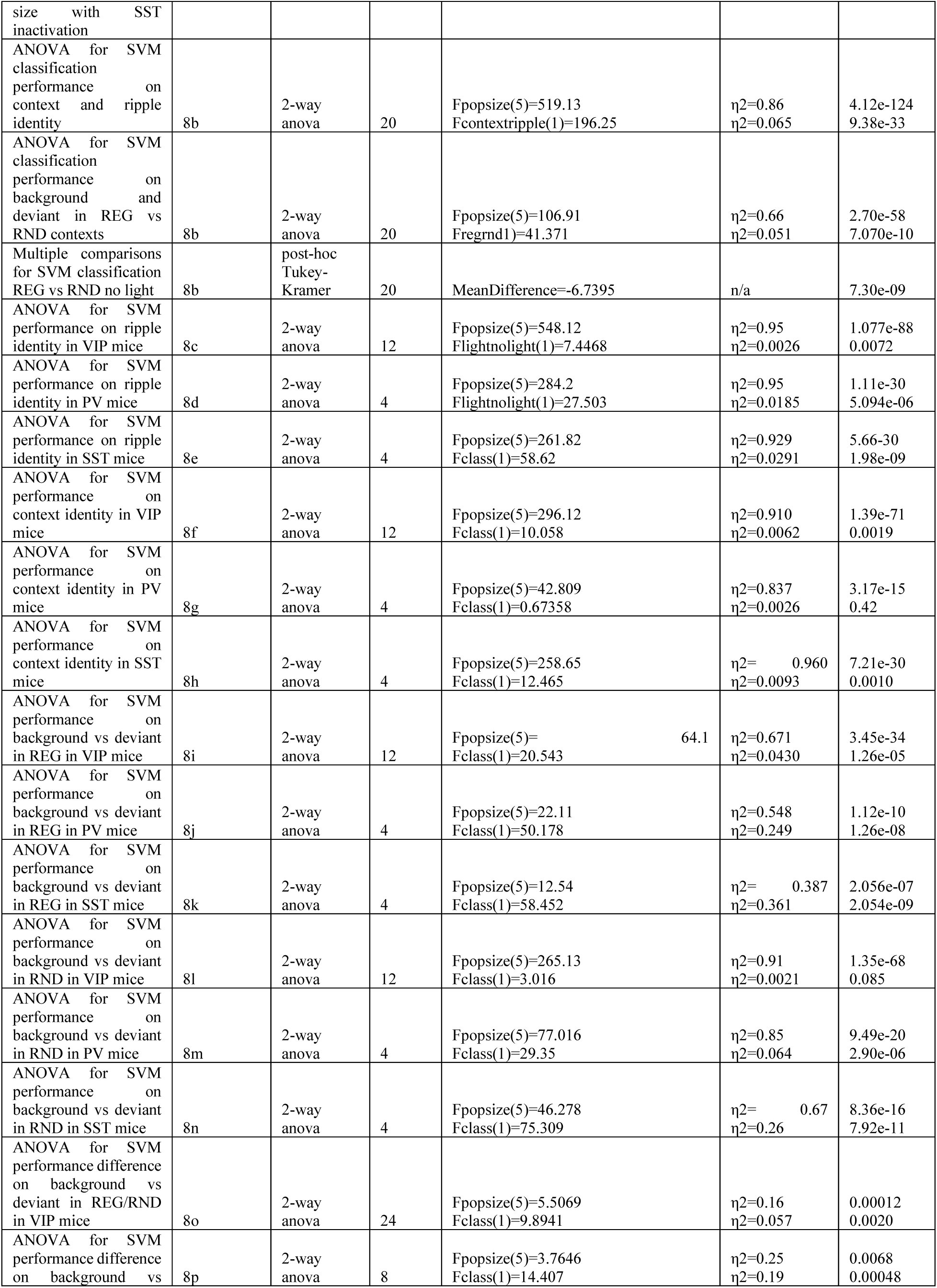
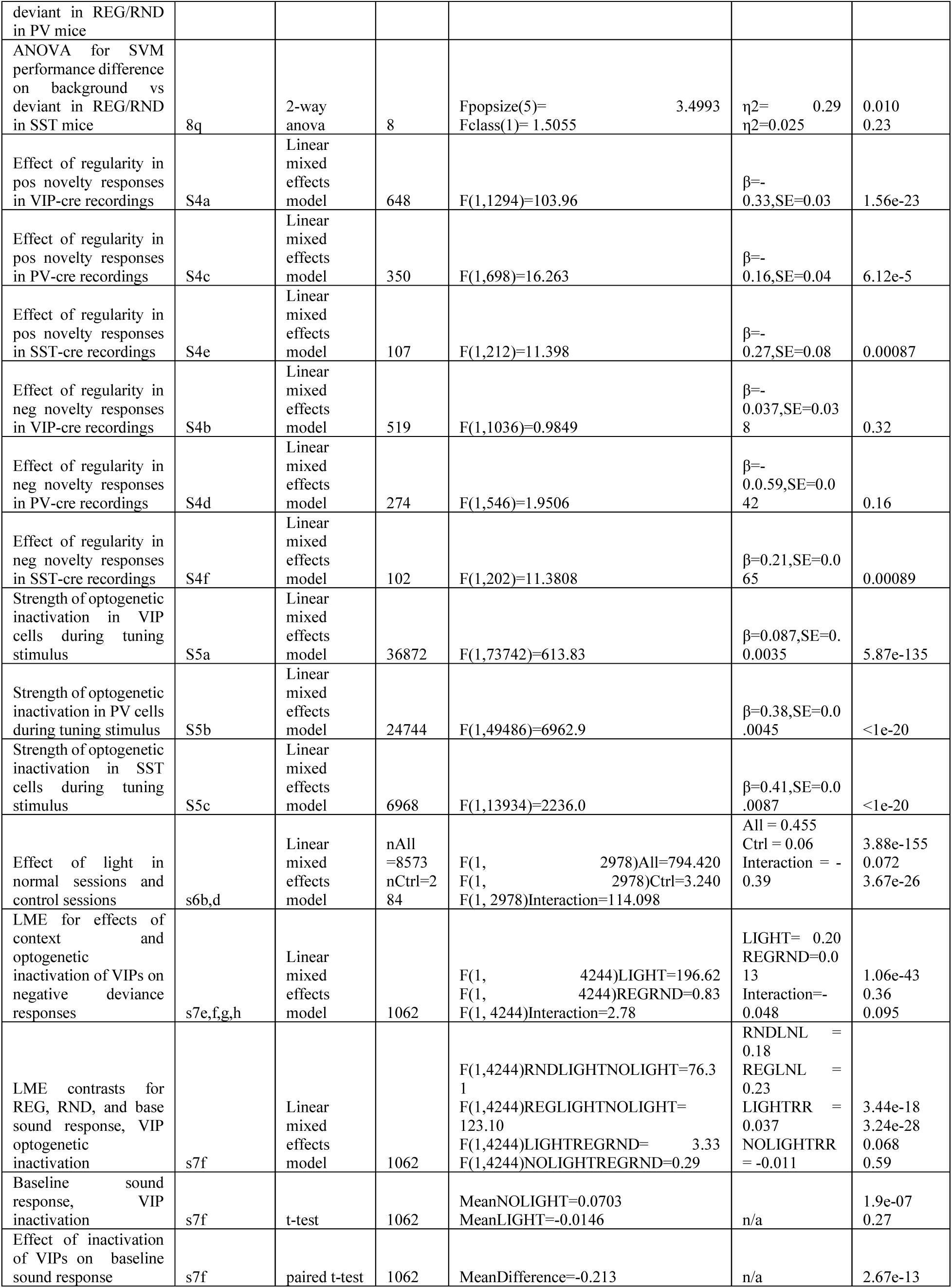
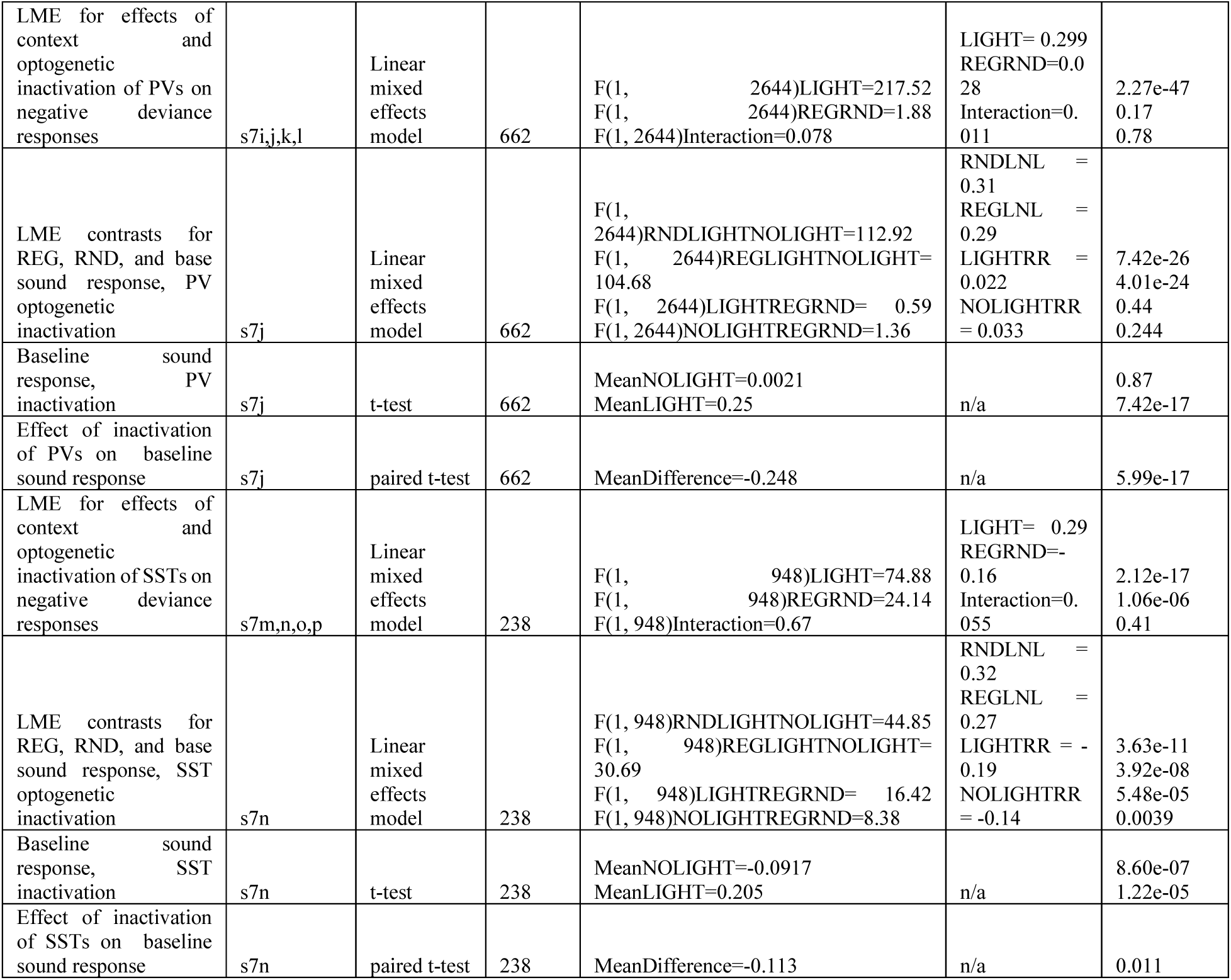

### Optogenetic Manipulation

Optogenetic stimulation was performed using a LED (Thorlabs M625L4) delivering 625 nm light through the objective lens (Tobin et al., 2024). Before each stimulus session, the LED light power was calibrated to ∼2.8 mW/mm^2^ for mice injected with the virus encoding Jaws from the Penn Vector Core and ∼7 mW/mm^2^ for mice injected with the virus encoding Jaws from the UNC Vector Core based on reported inhibition performance of Jaws (Chuong et al., 2014). Optogenetic manipulation occurred randomly in 50% of presentations of the deviant ripple. The LED was turned on continuously for 300 ms, synced with stimulus onset, during the deviant ripple presentation. Control mice (SST-Cre, n=2) were injected with the viruses encoding tdTomato (in a Cre-dependent fashion) and GCaMP, but not Jaws (Supplementary Figure S6), and did not exhibit a significant effect of light on tuning stimulus ripple responses (N =284, linear mixed effects model, F(1, 2978) = 3.24, p = 0.072).

### Acoustic stimuli

Auditory moving ripples were generated at a sampling rate of 400 kHz in MATLAB according to the following formula:

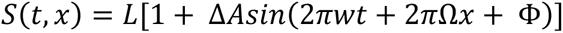

where *w* (cyc/s) is the temporal modulation parameter and Ω (cyc/octave) is the spectral modulation parameter. The variable Φ represents the random phase for the envelope, Δ*A* represents the sinusoidal amplitude, and L represents the depth modulation. The frequency is represented as 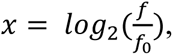 where f_0_ is the lowest frequency in the stimulus (Depireux et al., 2001). Individual ripples spanned a frequency range of 5-50 kHz, lasted 300 ms, were presented at 70 dB SPL with a 15 dB modulation depth, and had a 5 ms cosine ramp applied at the beginning and end. Background sequences were generated by concatenating ripples A, B, C, and D in either a fixed sequence (REG) or a random sequence (RND) resulting in a 1.2 s stimulus (Figure 1C). The deviant sequence was generated by concatenating ripples X, Y, Z, and W in that order for all stimulus conditions. Sequences were presented with a 1 s inter-stimulus interval. Background sequences were repeated 3-6 times pseudo-randomly before a deviant sequence was presented. A control sequence of each ripple presented in isolation with a 1 s ISI was also presented to determine the base sound response of neurons.

### Electrophysiology

Mice were prepared for *in vivo* electrophysiological recordings as described previously (Vogler et al., 2024). At least two weeks prior to recordings, a ground pin was inserted via small craniotomy over the left frontal cortex, and a custom-made stainless steel headplate (eMachine Shop) was secured to the skull using dental cement (C&B Metabond) and acrylic (Lang Dental) (Vogler et al., 2024). Recordings were performed in a custom-built acoustic isolation booth. At least one week following headplate surgery, mice were habituated to the recording setup for increasing durations (5-60 min) over the course of 3 days. On the day of recording, mice were anesthetized with 1.5-2.5% isoflurane, and a small craniotomy was made over the left auditory cortex using a dental drill, centered ∼1.5mm anterior to the lambdoid suture along the caudal end of the squamosal suture. Following the craniotomy, mice were allowed to recover from anesthesia for at least 30 min. Acute extracellular recordings in the auditory cortex were performed using a 32-channel silicon probe (A1x32-Poly2-5mm-50s-177, NeuroNexus). The probe was positioned perpendicularly to the brain surface and lowered at 1-2 μm/s to a final depth of 900-1200 μm or until all channels were within the brain. During lowering of the probe, we monitored responses to brief broadband noise clicks to confirm localization within auditory cortex. Neuronal signals were amplified and digitized with an Intan headstage (RHD 32ch) and recorded by an OpenEphys acquisition board (Siegle et al., 2017) at a rate of 30 kHz. Signals were filtered between 500 and 6000 Hz, offset corrected, and re-referenced to the median across all channels. Spiking data were sorted using Kilosort2 (Pachitariu et al., 2016) and manually curated in phy2 to identify putative single- and multi-units.

### Immunohistochemistry

Mouse brains were extracted and sliced as previously reported (Wood et al., 2022; Vogler et al., 2024). In brief, mice were anesthetized with an injection of dexmedetomidine (1 mg/kg) and ketamine (120 mg/kg) and then perfused with phosphate-buffered saline (PBS) and 4% paraformaldehyde (PFA). Brains were fixed in 4% PFA overnight at 4°C. Coronal slices were acquired on a cryostat (Leica CM1860) at −25°C. Slices were incubated overnight at 4°C with an RFP antibody (1:1000, Rockland: 600-401-379) followed by an anti-rabbit secondary for 2 hours at room temperature before imaging (see representative images from VIP-Cre, PV-Cre and SST-Cre mice of neurons labels with tdTomato and GCaMP in Supplementary Figure S1). Expression of GCaMP was identified and cross-referenced with the Paxinos mouse brain atlas (Figure 1A bottom).

### Data analysis and statistics

Suite2P (Pachitariu et al., 2017) was used to register raw two-photon images, select regions of interest (ROI), and perform neuropil correction (Betzel et al., 2019; Wood et al., 2022; Tobin et al., 2024). A 0.3 Hz highpass filter was used to correct for slow fluctuations in the calcium signal. The calcium signal from each ripple sequence was converted to a z-scored ΔF/F_std_=(F-mean(F_baseline_))/std(F_baseline_) where F_baseline_ is the calcium signal from the 300 ms preceding stimulus onset. An additional motion correction step was performed using PatchWarp (Hattori and Komiyama, 2022). We found that small motion artifacts during imaging induced negative transients in the calcium trace. These artifacts were identified by setting a threshold for the X and Y values from the summary stat output and any trials within 3 frames of an artifact were removed from further analysis. This removed approximately 10% of all recorded trials. For statistical comparison of responses, we computed the mean response strength during the 300 ms following the onset of a ripple on each trial. To identify neurons exhibiting significant deviance responses, we compared the responses over trials during the same ripple presented as deviant and tuning stimulus using an unpaired t-test in either random or regular context. To compare deviance response strength across neurons in different conditions, we computed mean response to the first ripple X in the deviant stimulus sequence and implemented repeated measures ANOVA with context (regular or random) and light (on or off) as two factors. We used Tukey-Kramer for post-hoc comparisons. For linear mixed effects models (LME), we used the fitlme function in MATLAB. Throughout our LME analyses, we included the recording session ID as a predictor to control for effects of session-to-session variability. If the output of the statistical test in MATLAB resulted in “p = 0”, this was most likely due to a rounding error, and we reported it as p < 1e-20. The results from all statistical tests are reported in Supplementary Table S1.

#### REGRND Index

The REGRND index was computed as the difference between the absolute values of the mean response to the deviant ripple presented in the REG and RND contexts divided by the sum of the absolute values. This provides a value between -1 and 1 where an index value close to 1 indicates that the neuron preferred to respond to the REG context, a value close to -1 indicates a preference for RND, and a value close to 0 indicates no preference.

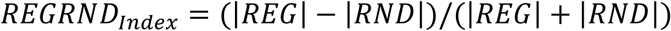

#### Exponential Fits

Mean responses *y* from 0-300 ms after stimulus onset for individual trials were fit to the exponential function below:

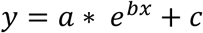

The variables *a, b,* and *c* were free parameters, and the variable *x* represented the trial presentation number. A neuron was considered to have a significant adaptation effect (*a > 0*) or facilitation effect (*a <* 0) if the RMSE from the exponential fit was less than the RMSE to the mean response. Fits were performed using MATLAB curve fitting toolbox.

#### SVM

Sample populations of 1, 12, 25, 50, 100, or 200 neurons were selected randomly (with replacement) from neurons within a single recording session. For each recording session, 1000 samples of responses were selected at random for each population size and average performance was reported. For each sample, the population response was represented as a vector with each entry being the mean response of a neuron 0-300 ms from stimulus onset for an individual trial. Then, a support vector machine (SVM) was trained to predict either ripple identity from the tuning stimulus or to predict the background context (REG/RND) from the responses to deviant ripple (Vapnik, 2013; Wood et al., 2022; Tobin et al., 2024). 20% of the trials were held out as a test set while the SVM was trained on the remaining 80%.

### Data and code availability

All original data and code will be made freely available upon publication. Source data will be deposited in a Dryad data depository. Data analysis code will be available on GitHub: https://github.com/geffenlab/Ding_2025.

## RESULTS

### Characterizing the sound response to individual moving ripples

We imaged calcium activity in populations of neurons in layer 2/3 of auditory cortex (AC) in awake, head-fixed mice presented with a series of sequences of spectro-temporally complex sounds (termed “moving ripples”) (Figure 1 A-D). Each ripple was comprised of a broadband sound with the envelope and frequency modulation varying in time. For the regular (REG) and random (RND) background ripples, we used the same 4 ripples (300 ms long, 5-50 kHz, 70 dB SPL mean intensity), which were repeated in the same sequence (REG; ABCD, ABCD, ABCD, …), or in varying order (RND; BADC, BDAC, CBDA, …). Each background sequence consisted of 3-6 repeats of the 4 ripples with 1 s inter-repeat interval followed by the deviant ripple sequence. Because the background sequence is comprised of the same ripples in both REG and RND, the only difference between the two contexts is the temporal order. The deviant sequence of ripples included 4 different ripples from those used in the background. These ripples were in the same frequency range but had different modulation frequencies. Additionally, the ripples were also presented individually with a 1 sec interstimulus interval to construct baseline tuning responses.

This stimulus allowed us to test multiple aspects of the neuronal responses to sounds in varying statistical regimes. First, we tested whether a given ripple evoked a greater or smaller neuronal response when it was unexpected. Second, we tested whether a deviant ripple elicited a greater or smaller response in the regular or random context. Third, we tested whether the responses to the context ripples differed when they were presented in regular or random context, and whether adaptation to their presentation over subsequent repeats differed based on whether their order was regular or random. We expected the responses to deviant ripples to be greater than the respective baseline sound response. We also expected deviance responses to be greater in the regular than random context. Finally, we expected adaptation to repeated ripple sequences to be stronger for regular than random contexts.

We recorded activity from 8,573 neurons in the AC, with 76.6% (n = 6,566) showing a significant sound response to at least one ripple (z-scored averaged fluorescent response 0-300 ms post stimulus-onset versus the average fluorescence 0-300 ms before stimulus on each trial, t-test with a Bonferroni correction for the number of ripples). The fluorescence of neurons typically increased over the first 200 ms after sound presentation followed by a gradual decrease in fluorescence (representative neurons, Figure 1E, F). Across all imaged neurons, 22.4% (n = 1,922/8,573) responded to only one ripple while 6.3% (n = 541/8,573) had significant responses to all eight ripples. Additionally, the fraction of responsive neurons to each ripple was relatively even across all eight ripples, with a maximum difference of 22.4% (Figure 1G, H, Supplementary Figure S3). Whereas many neurons exhibited an enhancement in fluorescence in response to the ripples (n = 1,967/6,566), the majority of imaged neurons exhibited a suppression in fluorescence to individual ripples (n = 4,599/6,566, Figure 1I). The suppression in fluorescence corresponds to a decrease in calcium concentration, which most likely is due to reduced spiking activity in response to the moving ripples (Attinger et al., 2017; Solyga and Keller, 2025). We then split the distribution of responses by mouse line (VIP-cre, PV-cre, and SST-cre) and found similar results. The majority of neurons from each line responded only to 1 ripple (VIP: 27.2%, PV: 31.3%, SST: 33.3%) and responses to ripples were relatively even across all eight ripples (maximum difference, VIP: 19.4%, PV: 26.1%, SST: 31.6%) with negative responses forming the majority (Supplementary Figure S3 A, B, C right). Furthermore, mean response strengths to tuning stimuli were similar across all three mouse lines (approximately -0.2 z-scored delta F/F, Supplementary Figure 7 A, B, C solid lines). When appropriate, we analyzed the positive responses as a separate group from negative responses. Combined, responses to individual ripples demonstrate that these stimuli efficiently drive neuronal calcium responses in AC.

### Regularity of context modulates deviance response

Most neurons in AC that we imaged exhibited a deviance response: 51.2% of neurons exhibited a stronger response to the deviant ripple in either the RND or REG condition as compared to the tuning response (significant change in mean response within the 300 ms window post-stimulus onset to the deviant ripple compared to tuning response to that ripple, N = 4,391 responsive neurons out of 8,573 neurons recorded). There was a significant effect of deviance on the response to the same ripple (n = 4,391, repeated measures ANOVA, f(2) = 51.18, p = 7.95e-23). These deviance responses were significant when computed separately for the positive (n = 2,429, repeated measures ANOVA, f(2) = 1816.91, p < 1e-20, Figure 2A-D) and negative (n = 1,962, repeated measures ANOVA, f(2) = 1324, p < 1e-20, Figure 2F, G, H) responding neurons. The remaining neurons (n = 4,182) exhibited no significant difference in response to the deviant ripple compared to its baseline response (Figure 2J, K, L). Deviance responses were also significant for a group that included neurons with both positive deviance and baseline responses (n = 191, repeated measures ANOVA, f(2) = 129.72, p = 1.14e-43, Figure 2N, O, P).

**Figure 2.**
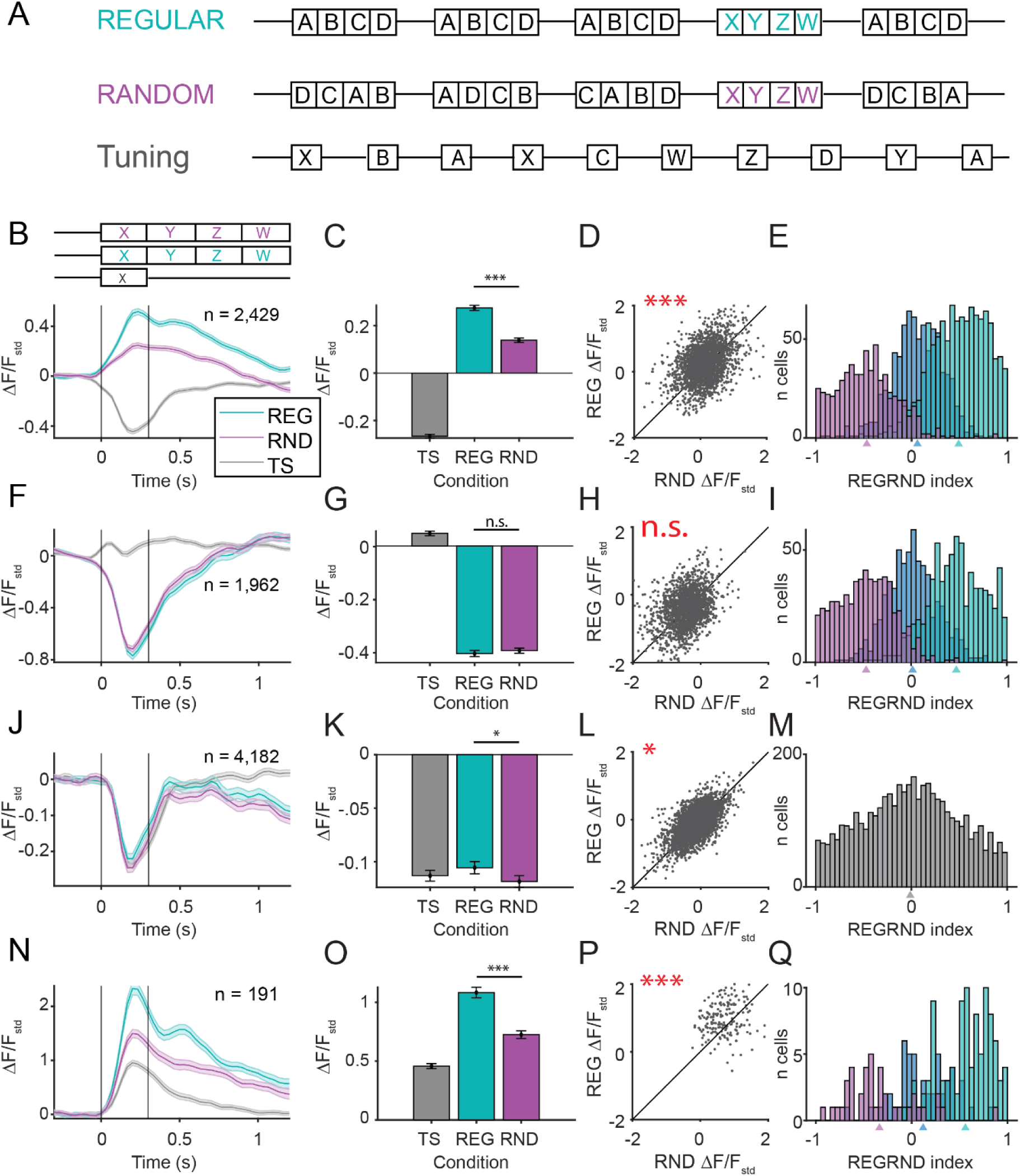
AC neurons exhibit stronger deviance responses in regular than random contexts. **A.** Schematic of the stimuli for REG and RND contexts (sequence of 4 ripples ABCD) with deviant ripples (XYZW). A tuning stimulus comprising each of 8 ripples was used to identify the baseline sound response to the ripples for individual neurons. **B.** Population-averaged calcium traces across neurons with positive deviance response to the deviant ripple presented in the REG context (cyan), RND context (magenta), and as part of tuning stimulus (grey) as z-scored ΔF/F on the left. Vertical lines represent duration of the first deviant ripple. **C.** Mean responses to the deviant ripple over 300 ms post stimulus onset in REG, RND and tuning (TS) contexts. **D.** Scatter plot of responses to deviant ripple under REG and RND contexts. **E.** Histogram of the REGRND index for responses to the deviant ripple for neurons with preference for REG context (cyan), RND context (magenta) or no preferences (gray). Mean values for the index for each group are denoted by triangles below the x-axis. **B, C, D, E.** Responses of neurons that exhibit positive responses to deviant ripple under either REG or RND contexts. The mean responses show a significant effect of context on the response amplitude (n = 2,429, lme, p < 1e-20) with deviance responses in REG context being the strongest (REG), followed by deviance responses in RND context (RND), and then the tuning stimulus responses (TS) (post-hoc Tukey-Kramer, p = 5.96e-08). **F, G, H, I.** Similar to B-E, but for responses of neurons that exhibit negative responses to deviant ripples under either REG or RND context (n = 1,962). For negative responses, there is no difference between REG and RND deviance responses (lme, n.s.). **J, K, L, M.** Similar to above, but for all neurons without a significant deviance response (n = 4,182, lme, n.s.). **N, O, P, Q.** Similar to above, for all neurons with both a positive sound response and a positive deviance response (n = 191, lme, p = 4.11e-15). REG responses are the strongest, followed by RND responses, followed by TS responses (post-hoc Tukey-Kramer, p = 5.96e-8).

For neurons with positive deviance responses, the REG context enhanced the deviance response in the corresponding direction. The deviance response magnitude was greater under the REG context than under the RND context (lme: F(1, 7284) = 121.761, p< 1e-20, Figure 2B, C, D). The context-dependent difference in deviance response strength persisted when we considered neurons with both positive deviance and baseline sound responses to the deviant ripple (lme: F(1, 570) = 65.161, 4.11E-15, Figure 2N, O, P). For negative-responding neurons, the difference in deviance response between the REG and RND contexts was not significantly different (lme: F(1, 5883) = 0.719, p = 0.4, Figure 2G, H). Similarly, neurons that did not exhibit significant deviance responses did not have a preference for the REG context (lme: F(1, 12543) = 2.965, p = .085, Figure 2J, K, L). For positive deviance responses, these results are consistent with our predictions, as we expected the deviance response to be stronger in the regular compared to random conditions. We note that a substantial fraction of neurons exhibited negative calcium responses to sound stimulation, with a distribution of responses similar across the VIP-Cre, PV-Cre and SST-Cre lines (Supplementary Figure S2). Similar suppressive signals in auditory cortex have been documented by other groups (e.g., Solyga and Keller, 2025; Tsukano et al., 2025), suggesting this is a reproducible phenomenon, but it is difficult to interpret negative novelty responses as a larger negative response may be a combination of a stronger or weaker interactions. Importantly, our key effects—including context-dependent deviance responses—were robust across different subsamples (Figure 2).

To further account for possible differences in responses by mouse line as well as any potential bias from our selection criteria, we separately analyzed our data from VIP-cre, PV-cre, and SST-cre mice (Supplementary Figure S4). We first selected neurons that showed a significant difference between REG and RND deviance responses and then split into populations with a positive or negative deviance response. For positive deviance response, we found that REG responses were significantly greater than RND responses across all three mouse lines (LME, VIP-Cre: N = 648, p< 1e-20, PV-Cre: 350, p=6.12e-5, SST-Cre: N = 107, p=.0009; Supplementary Figure S4 A, C, E). For only negative responses, we did not observe consistent differences across the three mouse lines, and context had varying effect on negative deviance (LME, VIP-Cre: N = 519, p = .32, PV-Cre: N = 274, p = .16; SST-Cre: N = 102, p = .0009; Supplementary Figure S4B, D, F). The effect in positive novelty responses is consistent with our hypothesis and results across all mouse lines.

To further understand the preference of individual neurons’ deviance responses for temporal context, we computed a REGRND index (see Methods) which ranged from -1 to 1, with -1 indicating a preference for the RND context and 1 indicating a preference for the REG context. For positive deviance responses, we found that a majority of neurons only exhibited a significant deviance response in one of the REG (43.8%) or the RND (24.5%) contexts. For neurons exhibiting a significant deviance response in both contexts, we found a significant preference for the REG context when considering the REGRND index (mean = 0.06, t-test, p = 1.31e-10, Figure 2E). This preference for the REG context was stronger when we only considered neurons with a positive deviance response and a positive baseline sound response (mean = 0.12, t-test, p =0.0046, Figure 2Q). On the other hand, neurons with negative deviance responses exhibited a small difference in the percentage of neurons selective for the REG (36.9%) and RND (30.8%) contexts and no significant difference in the REGRND index for neurons responsive in both (mean = 0.015, t-test, p = 0.14, Figure 2I). Thus, positive deviance responses increase in magnitude as well as in the fraction of responsive neurons in the REG over RND contexts, whereas negative deviance responses have mixed context-dependent effects.

We next analyzed how the deviance response changes over the course of an entire REG or RND stimulus block. Previous work has shown that adaptation in AC occurs over multiple time scales (Ulanovsky et al., 2004). We hypothesized that temporal regularity may also impact adaptation on longer time scales in a similar manner, with stronger adaptation occurring in the REG context compared to the RND context. We fit an exponential response function to the time course of per trial responses from individual neurons (Figure 3A, B, E). We found that a sizeable fraction of neurons exhibited long term adaptation (48.03%) or facilitation (41.52%) to the deviant ripple presentation. Neurons adapted or facilitated faster in the REG as compared to RND context (Figure 3C, D, F, G): time constants for both adaptation and facilitation were greater in the RND context than the REG context (n_adapt REG_ = 2,545, n_facil REG_ = 2,032, n_adapt RND_ = 2,224, n_facil RND_ = 1,843, 2-way ANOVA, f(1) = 10.29, p = 0.0044), indicating that context has a secondary effect on the long-term adaptation of neuronal responses to the deviant ripple. These results demonstrate a secondary mechanism for adaptation to stimulus context during deviance detection.

**Figure 3.**
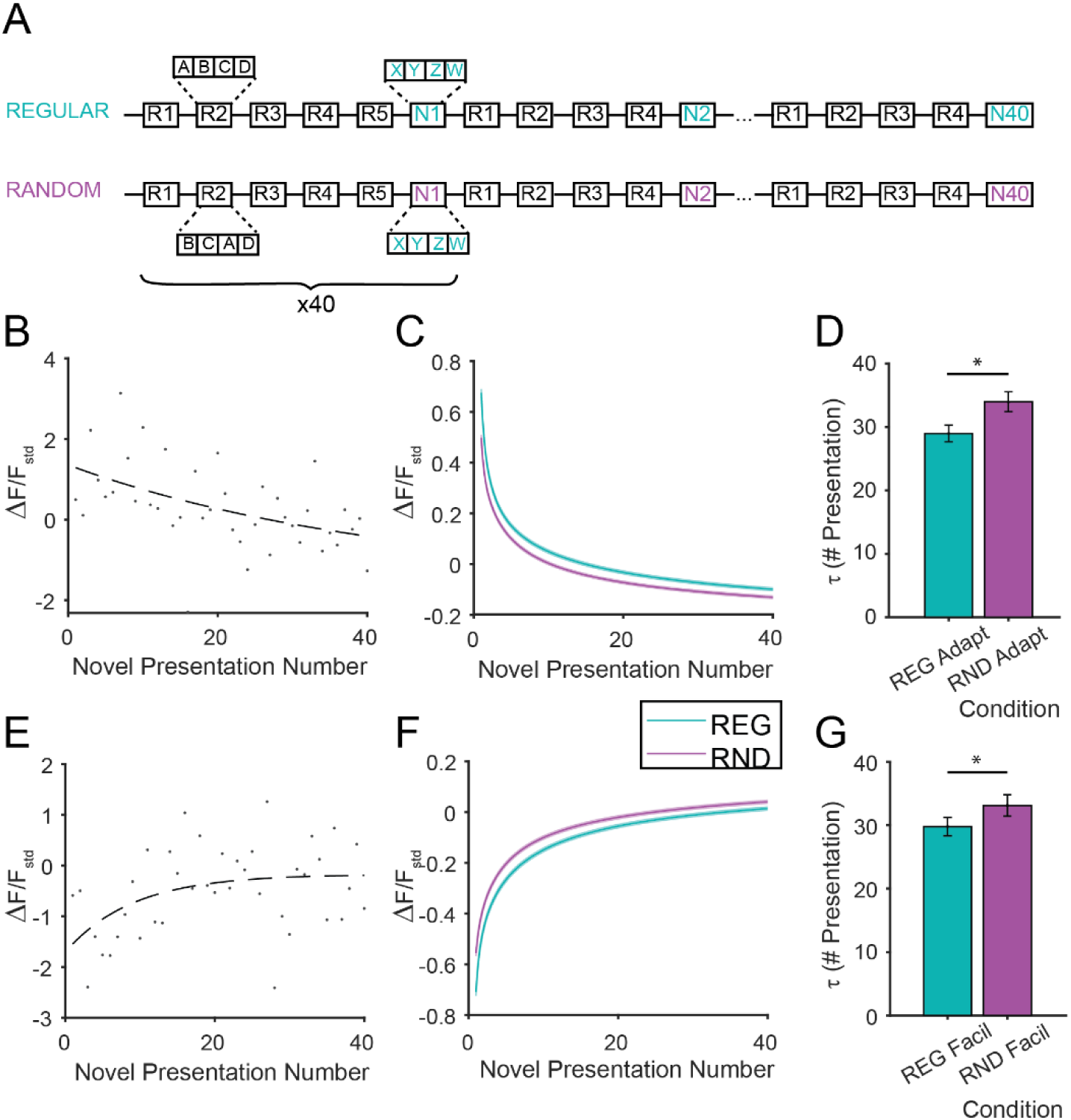
Temporal regularity modulates long-term deviance response dynamics. **A.** Ripple sequences were presented in blocks of entire REG or RND contexts. The deviant ripple is always the same, allowing for potential adaptation to the deviant ripple itself. **B, E.** Deviance responses over successive deviant stimulus presentations, for example cells exhibiting adaptation (B) and facilitation (E) in neuronal responses to the deviant ripple sequence in a REG context. Dots denote response strength for a individual deviant ripple presentation and the dashed line is the best exponential fit. **C, F.** Exponential fits for adaptation (C) and facilitation (F) in REG (cyan) and RND (magenta) contexts across the population. **D, G.** Mean time constants for adaptation (D) and facilitation (G) in REG and RND contexts for all neurons that exhibit such long-term dynamics. Both adaptation and facilitation are faster in the REG context (lower time constant, 2-way ANOVA, p = 0.0044).

### Cortical neurons exhibit differential spiking responses to deviant ripples in regular and random contexts

To resolve finer temporal information in cortical responses, we next performed electrophysiological recordings in the auditory cortex of awake mice (n = 5) while presenting the ripple stimuli (Figure 4A-C). From these recordings, we identified deviance responsive units using a paired t-test between the spike rates from the 300 ms following stimulus onset when the ripple was presented in isolation versus as a deviant and found that most neurons exhibited a deviance response in either context (55%, n = 164 neurons). Among the units with a significant deviance response, 59% responded with an increase in firing rate (n = 96 neurons). When comparing the firing rates in response to the deviant ripple in the REG and RND contexts, there was a significant effect of context in the first 100 ms of the response (Figure 4D, E, repeated measures ANOVA, f(2) = 9.36, p = 0.00011). Deviance responses in the REG context were significantly greater than those in the RND context (post-hoc Tukey-Kramer, p = 0.024). Furthermore, the differential effect of context extended to the first 100 ms of the second (paired t-test, p = 0.017) and third (paired t-test, p = 0.019) ripples in the deviant sequence but not the fourth (paired t-test, p = 0.093). When examining the last 100 ms of the response to the first ripple, although the mean firing rate was higher in the REG context (∼10.2 Hz) compared to the RND context (∼9.2 Hz), there was no significant effect of context (repeated measures ANOVA, f(2) = 2.022, p = 0.13). These results indicate that the information about context is integrated early in the responses of neurons in AC and support our findings from 2-photon imaging.

**Figure 4.**
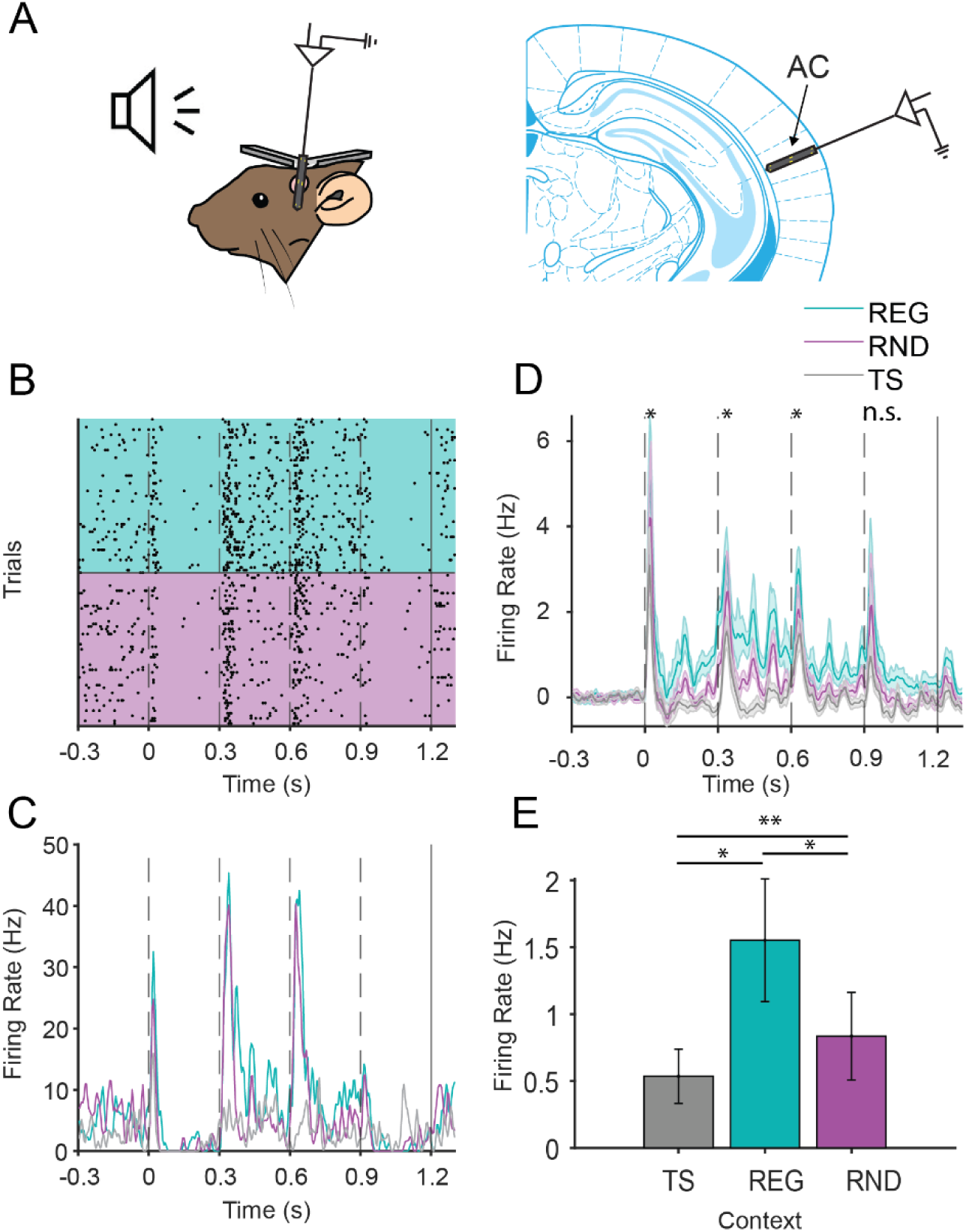
AC neurons exhibit higher firing rates to deviant ripples in regular as compared to random contexts. **A.** Left: schematic of electrophysiological recordings from AC in awake head-fixed mice. Right: Electrode superimposed on the AC outline from the Paxinos Brain Atlas (Paxinos and Franklin, 2012). **B.** Raster plot of spike times from an example neuron in response to the deviant ripple sequence in REG (cyan) and RND (magenta) context. **C.** PSTH of the same neuron’s firing rate in response to the deviant ripple sequence in REG and RND as well as the tuning response to the first ripple in the deviant sequence (grey). **D.** Mean firing rates over all neurons with a significant deviance response (n = 164) to the deviant ripple sequence in REG (cyan), RND (magenta), and to the first ripple in isolation (grey). **E.** Mean firing rates in the first 100 ms post stimulus onset in REG, RND, and tuning contexts show a significant effect of context on deviant ripple responses (repeated measures ANOVA, f(2) = 9.36, p = 0.00011). Error bars represent SEM.

### Cortical inhibitory neurons exhibit contextually modulated deviance responses

Inhibitory neurons modulate deviance responses in AC (Natan et al., 2015; Yarden et al., 2022) as well as other sensory areas (Millman et al., 2020; Furutachi et al., 2024). In AC, cortical inhibitory neurons also exhibit the enhanced deviance response associated with SSA (Natan et al., 2015). We investigated whether cortical inhibitory neurons exhibit a similar deviance response to our ripple stimulus by imaging from a VGAT-Cre mouse line crossed with a transgenic Cre-dependent GCaMP8m line. This crossed line results in GCaMP8m expression only in inhibitory neurons. We found that, similar to our results from our pan-neuronal recordings, inhibitory neurons also exhibit a context-modulated deviance response (Figure 5). We found that 74.3% of cortical inhibitory neurons exhibited a significant deviance response. The REG context enhanced the response compared to the RND context for positive responses (lme: F(1, 2592) = 152.299, p < 1e-20, Figure 5C-E), and negative responses in the REG context were slightly stronger than in the RND (lme: F(1, 3426) = 3.99, p = .046, Figure 5G-I). In a subset of recording sessions, we presented a control stimulus sequence where the ripples used (ABCD or XYZW) for the background and deviant sequences were swapped. Neurons from these sessions also exhibited an enhanced deviance response in the REG context compared to the RND context (lme: F(1, 771) = 112.939, p < 1 e-20, Figure 5O-Q). The REGRND index from the inhibitory neurons followed a similar pattern to our results from the general neuronal recordings. For positive responses, there is a significant preference for the REG context in neurons responsive to both contexts in the original (REGRND index mean = 0.098, t-test, p = 1.31e-10, Figure 5F) and control sequences (REGRND index mean = 0.154, t-test, p =7.02e-9, Figure 5R), while no significant effect was observed for negative responses (REGRND index mean = 0.012, t-test, p = 0.25, Figure 5J). Furthermore, the inhibitory neuronal population that did not exhibit a significant deviance response had a slightly stronger novelty sound response in regular condition, although this relation was not significant in a post-hoc test (lme: F(1, 2085) = 6.537, p = 0.011, post-hoc Tukey-Kramer, n.s., Figure 5K-N). The large fraction of context-sensitive inhibitory neurons and strong context-evoked differences in the deviance responses in inhibitory neurons indicate that the deviance response found in the general population may be controlled by inhibitory neurons.

**Figure 5.**
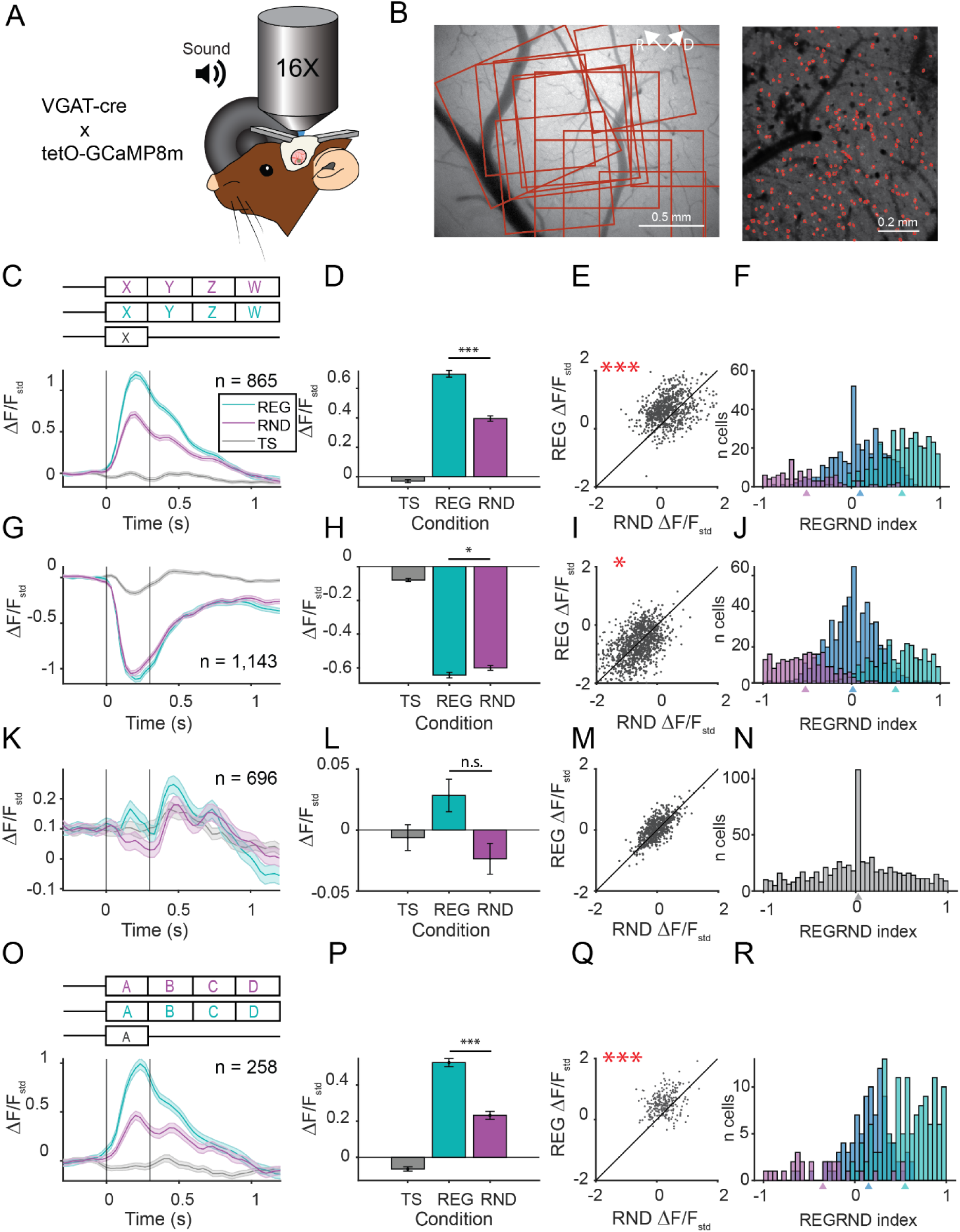
Inhibitory neurons in AC exhibit context-dependent deviance responses. **A.** Schematic of 2-photon imaging from a transgenic mouse line expressing GCaMP8m only in inhibitory neurons. **B.** Left: Representative fields of view imaged (white boxes) overlayed on AC. For clarity, imaging areas close to one another have been combined. Right: representative field of view with identified neurons outlines using Suite2p. **C, G, K, O.** Population-averaged calcium traces across neurons with positive deviance response to the deviant ripple presented in the REG context (cyan), RND context (magenta), and as part of tuning stimulus (TS, grey) as z-scored ΔF/F on the left. Vertical lines represent duration of the first deviant ripple. **D, H, L, P.** Mean responses to the deviant ripple over 300 ms post stimulus onset in REG, RND and TS contexts. **E, I, M, Q.** Scatter plot of responses to the deviant ripple under REG and RND contexts. **F, J, N, R.** Histogram of the REGRND index for responses to the deviant ripple for neurons with preference for REG context (cyan), RND context (magenta) or no preferences (gray). Mean values for the index for each group are denoted by triangles below the x-axis. **C, D, E, F.** Responses to the deviant ripple over neurons with positive deviance responses (n = 865). REG deviance responses are significantly greater than RND deviance responses (lme, p < 1e-20). **G, H, I, J.** Similar to C-F, but for all negative deviance responses (n = 1,143). For negative responses, REG deviance responses were lower than RND responses (lme, p= 0.046). **K, L, M, N.** Similar to C-F, but for average responses of all neurons without a significant deviance response (n = 696). There is an increase in deviant responses in REG as compared to RND context (lme, p = 0.011). However, this was not significant in post-hoc Tukey-Kramer test. **O, P, Q, R.** Similar to C-F, but for recordings to stimuli in which background and deviant ripples were swapped (ABCD was the deviance sequence and XYZW was the background sequence) (n = 258, lme, p < 1e-20). Responses in REG context are the strongest, followed by responses in RND context, followed by TS responses (post-hoc Tukey-Kramer, p = 1.19e-07).

### Optogenetic inactivation of inhibitory neurons differentially modulates deviance responses

We next asked how specific classes of inhibitory neurons modulate cortical deviance detection in different contexts. Previous work has shown that PV and SST neurons differentially modulate stimulus-specific and temporal adaptation in AC (Natan et al., 2015, 2017). VIP neurons form a disinhibitory circuit with SST neurons and have also been shown to modulate deviance responses in sensory cortex (Furutachi et al., 2024). Therefore, we hypothesized that these distinct inhibitory neuron classes differentially modulate deviance responses in regular and random contexts. To investigate the role of inhibitory neurons in modulating the deviance response, we measured calcium activity in AC neurons while inactivating PV, SST, or VIP neurons using optogenetic activation of Jaws on a subset of trials (Figure 6A-D). Jaws is a red-shifted inhibitory opsin which we expressed selectively in each of three interneuron subtypes using Cre-expressing transgenic mice. We inactivated each of the three inhibitory neuron classes during the presentation of the first ripple in the deviant sequence to identify their selective role in modulating the response. As expected, inactivating VIP neurons decreased response amplitude to ripples, whereas inactivating SST or PV neurons increased the responses to ripples (Figure 6E-P, Figure S5, solid and dashed lines).

**Figure 6.**
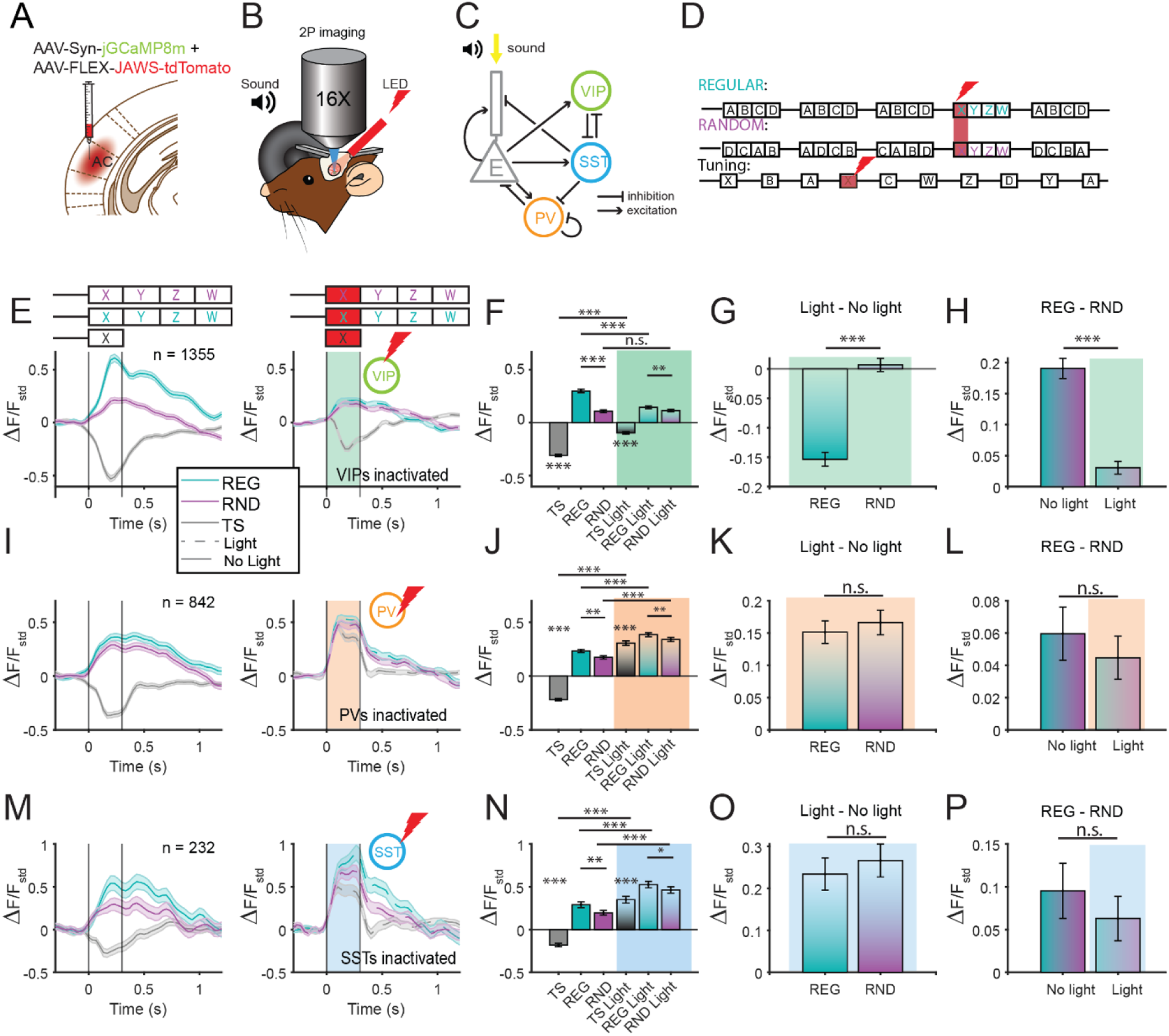
Selective context-dependent modulation of deviance responses by VIP neurons. **A, B, C, D.** Schematics of the experimental set-up. **A.** Viral injection targeted to A1 included injection of a virus encoding GCaMP and a virus encoding a JAW-tdTomato construct in a Cre-dependent fashion into VIP-Cre, PV-Cre, or SST-Cre mice. Image from the Paxinos Brain atlas (Paxinos and Franklin, 2012). **B.** 2P imaging was performed in mice injected with JAWS-tdTomato in addition to GCaMP. **C.** Simplified diagram of the inhibitory-excitatory microcircuit in A1 that includes PV, SST and VIP neurons. PV neurons largely target the excitatory neuronal cell bodies, SST neurons target distal dendrites, whereas the VIP neurons inhibit SST neurons. **D.** Diagram of the stimulus with the light delivered during the deviant stimulus presentation on a subset of trials. Optogenetic inactivation was performed during the first ripple in the deviant sequence. **E, I, M.** Population-averaged calcium traces across neurons with positive deviance responses to the deviant ripple presented in the REG context (cyan), RND context (magenta), and as part of tuning stimulus (grey) as z-scored ΔF/F. Left: light off; Right: light on. Vertical lines represent duration of the first deviant ripple. **F, J, N.** Mean responses to deviant ripple over 300 ms post stimulus onset in REG, RND and TS contexts. Left: light off. Right: Light on. **G, K, O.** Mean changes in deviance responses due to inactivation of inhibitory neurons under REG (cyan) and RND (magenta) contexts. **H, L, P.** Mean difference in deviance responses between REG and RND contexts without light (left) and with light (during inhibitory neuron inactivation, right). **E, F, G, H.** Effects of inactivation of VIP inhibitory neurons. Inactivation of VIP neurons reduces the deviance response (n = 1,355, linear mixed effects model, p = 5.02e-19). This modulation of the deviance response is context-specific, with a much larger reduction in deviance responses under REG as compared to RND context (linear mixed effects model, interaction term, p = 4.48e-11). **I, J, K, L.** Effects of inactivation of PV inhibitory neurons. Inactivation of PV neurons increases the deviance response (n = 842, linear mixed effects model, p = 2.86e-11) and does not result in a differential effect of context (linear mixed effects model, interaction term, p = 0.64). **M, N, O, P.** Effects of inactivation of SST inhibitory neurons. Optogenetic inactivation of SST cells also increases the deviance response (n = 232, linear mixed effects model, p = 1.1e-6) but does not result in a significant differential effect of context (linear mixed effects model, interaction term, p = 0. 63).

For neurons with a positive deviance response, inactivation of VIP neurons decreased the amplitude of the tuning response to the ripples (paired t-test, p < 1e-20, Figure 6E, F). The effect of VIP neurons on deviance responses was selective for the temporal context (linear mixed effects model, F(1,5416) = 43.57, p= 4.48e-11, Figure 6E-H): suppressing VIP neurons significantly reduced the deviance response in the regular (F(1,5416) = 80.01, p < 5.02e-19), but not random contexts (F(1,5416) = 0.15, p = 0.70, Figure 6F, G). Inactivating PV or SST neurons increased tuning responses to the ripple, changing the mean response from negative to positive (paired t-test, p_PV_ < 1e-20, p_SST_ < 4.19e-29, Figure 6I, J, M, N). Inactivating PV or SST neurons increased the deviance response for both REG (F_PV_(1, 3364) = 44.57, p_PV_ =2.86e-11, Figure 6I-L, F_SST_ (1, 924) = 24.07, p_SST_ = 1.1e-6, Figure 6M-P) and RND (F_PV_(1, 3364) = 53.76, p_PV_ =2.82e-13, Figure 6I-L, F_SST_ (1, 924) = 31.18, p_SST_= 3.1e-8, Figure 6M-P) contexts. However, there was no difference in the increase in deviance response due to the temporal context (linear mixed effects model, F_PV_ (1,3364) = 0.22, p_PV_ (Context x Light) = 0. 64, Figure 6K, L, F_SST_(1,924) = 0.32, p_SST_ (Context x Light) = 0.63, Figure 6O, P). We also fit a linear mixed effects model to test whether the observed difference in context dependent light modulation was significantly across mouse lines. Inactivation of VIPs was the only significant context dependent effect in this model (linear mixed effects model, F_VIP_(1,2426) = 31.95, p_VIP_ = 1.76e-8; F_PV_(1,2426) = 0.339, p_PV_ = 0.56; F_SST_(1,2426) = 0. 653, p_SST_ = 0. 42). Furthermore, the difference in effect between VIP and PV (linear mixed effects model, F(1,2426) = 7.13, p = 0.0077) and VIP and SST (linear mixed effects model, F(1,2426) = 7.95, p = 0.0049) was significant while the difference between PV and SST was not (linear mixed effects model, F(1,2426) = 0.083, p = 0. 77). Thus, VIP neurons selectively control the deviance responses in REG, but not RND context, whereas PV and SST neurons reduce deviance responses in a context-independent fashion.

For neurons that have negative deviance responses, optogenetic inactivation of all three interneuron classes modulated the activity during the deviant stimulus (Supplementary Figure S7). For negative deviance responses, inactivating VIP neurons decreased the amplitude of deviance responses under both random and regular contexts (linear mixed effects model, F(1,4244) = 196.62, p = 1.06e-43, Supplementary Figure S7E-H). This modulation of deviance responses was not significantly context-dependent (linear mixed effects model, F(1,4244) = 2.78, p(Context x Light) = 0. 095, Supplementary Figure S7G, H). Inactivating PV and SST neurons increased the deviance responses under both regular and random contexts (linear mixed effects model, F_PV_(1,2644) = 217.523, p_PV_ = 2.27e-47; F_SST_(1,948) = 74.88, p_SST_ = 2.12e-17, Supplementary Figure S7I-P). This change in deviance response was not context-dependent for either PV or SST neuron inactivation (PVs: F_PV_(1,2644) = 0.078, p = 0.78; SSTs: F_SST_(1,948) = 0.67, p = 0.41, Supplementary Figure S7K, L, O, P).These results suggests that the differential function of VIPs may be limited to positive novelty responses.

To understand how, across the positive and negative deviance responses, the inhibitory neurons affected deviance detection, we trained a support vector machine (SVM) classifier to differentiate between responses to a deviant ripple and a context ripple (Figure 7A). For each recording session, we trained the classifier on the responses to the deviant ripple X versus responses to the first ripple in the repeated background sequence. To match the position in the sequence as much as possible, we used the last repeat prior to the deviant ripple on each trial. For regular contexts, this was always A, whereas for random contexts, this ripple varied between A, B, C and D. We compared the performance of the classifier in predicting whether neuronal responses corresponded to the deviant or context ripple for trials with light (when the inhibitory neurons were suppressed) and without light, for the three different inhibitory neuron types.

**Figure 7.**
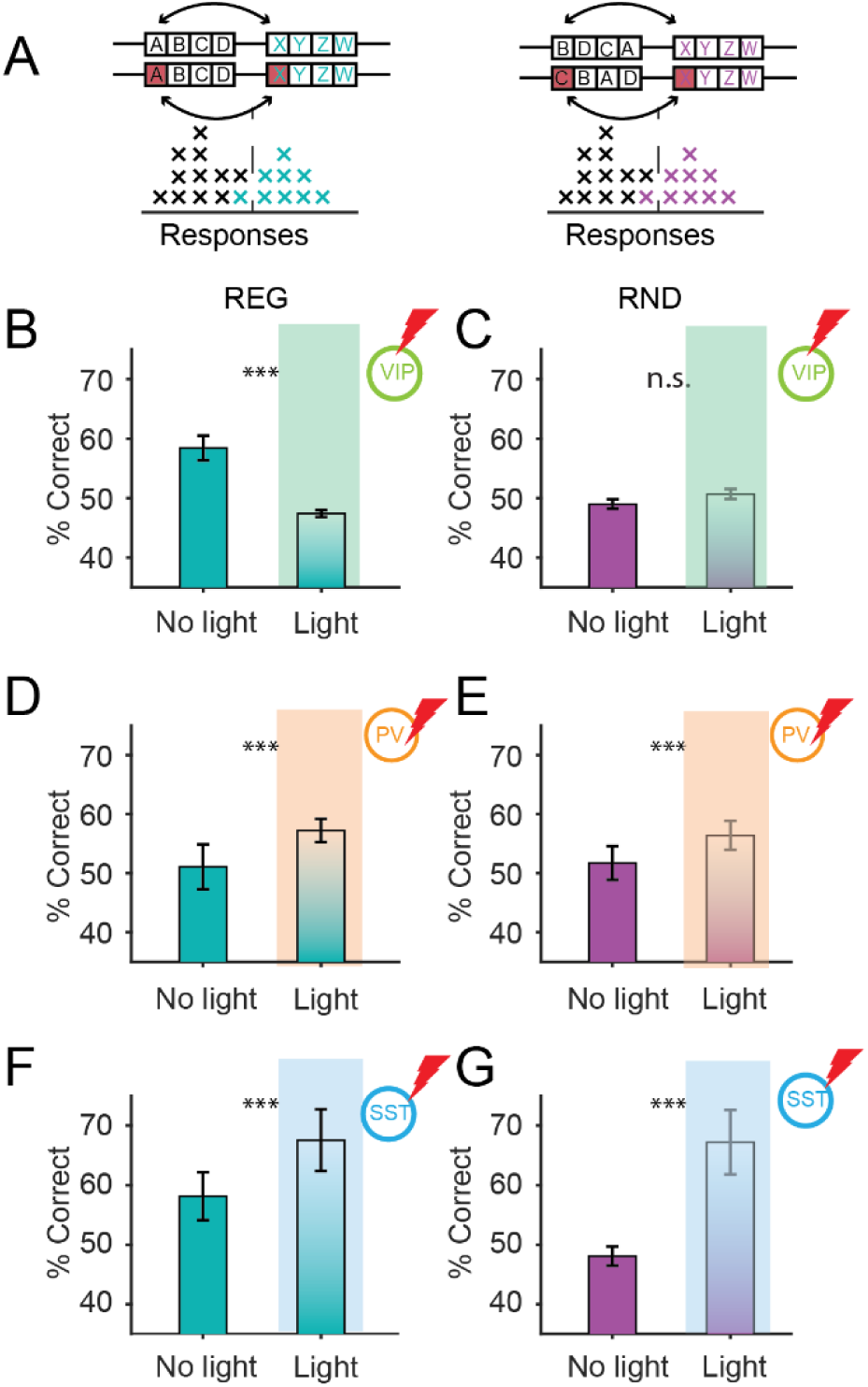
Differential effect of inhibitory neuronal inactivation on deviance detection under random and regular contexts. **A.** Schematic of SVM classification for single neuronal responses to deviant and context ripples. The activity during individual presentations of the context ripple and the deviant ripple X was used to train the classifier under REG and RND contexts and during light and no light trials. **B-G.** Single neuron SVM performance for trials with (left) and without (right) light in the REG (B, D, F) and RND (C, E, G) contexts. **B, C.** VIP inactivation decreases SVM performance in REG context (post-hoc Tukey-Kramer, p = 0.00036), but not in RND context (post-hoc Tukey-Kramer, p = 0.85). **D, E.** PV inactivation improves SVM performance in REG context (post-hoc Tukey-Kramer, p < 7.76e-7) and in RND context (post-hoc Tukey-Kramer, p = 0.00017). **F, G.** SST inactivation improves SVM performance in REG context (post-hoc Tukey-Kramer, p < 1.28e-7) and in RND context (post-hoc Tukey-Kramer, p < 4.98e-9).

We found that inactivating VIP neurons differentially modulated the classifier performance, reducing it only for the regular context condition, but not for the random context (post-hoc Tukey-Kramer, p_REG_ < 0.00036, post-hoc Tukey-Kramer, p_RND_ < 0.85, Figure 7B, C). In contrast, inactivating PV or SST neurons improved SVM performance under both random (post-hoc Tukey-Kramer, p_PV_ < 0.00017, post-hoc Tukey-Kramer, p_SST_ < 4.98e-9, Figure 7E, G) and regular (post-hoc Tukey-Kramer, p_PV_ < 7.76e-7, post-hoc Tukey-Kramer, p_SST_ < 1.28e-7, Figure 7D, F) contexts. These results are in line with the changes in the mean response strength that we observed (Figure 6). They furthermore suggest that the selective effects of inactivating VIP neurons for REG context persist for the discriminability of deviant stimuli, whereas PV and SST neurons affect deviance responses, but not in a context-independent fashion.

We next asked more broadly how these changes in deviance detection are implemented across the population of AC neurons. We expanded the discriminability analysis to multiple discrimination schemes and used subsets of progressively increasing numbers of neurons to train the SVMs. We found that discriminability increased with the number of neurons in the analysis and did not saturate at 200 neurons for any discrimination scheme (Figure 8A, B).

**Figure 8.**
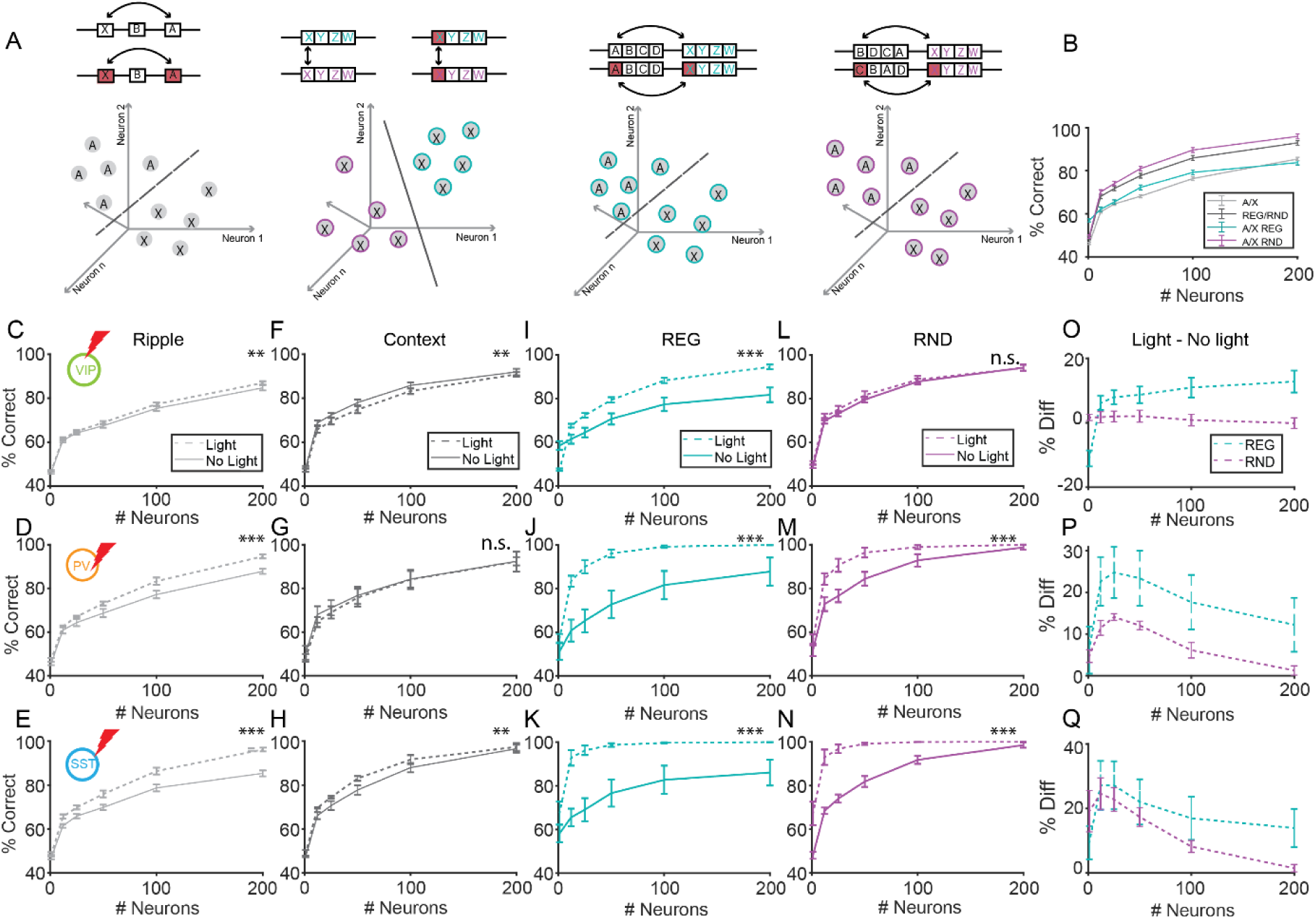
Deviance decoding is improved with inhibitory neuron inactivation at the population level. **A.** Schematic of a SVM identifying a linear decision boundary between points in a high dimensional space. We computed the SVM to discriminate between ripples A and X from the tuning response stimulus; for the deviant ripple X under REG and RND contexts; for the deviant ripple X versus background context ripple under REG; and RND contexts. All SVMs were trained and tested on data with and without optogenetic inactivation for each of the three inhibitory neuronal classes. **B.** SVM discrimination performance between context ripple and deviant ripple X from the sound response stimulus (grey); discrimination performance of the deviant ripple X between REG and RND contexts using the deviance response (black); deviance detection of X under REG (cyan) and RND (magenta) contexts. Context is easier to classify than ripple identity (2-way ANOVA, f(1)=196.25, p=9.38e-33). **C, D, E.** Effect of optogenetic inactivation of VIP (C), PV (D), and SST (E) neurons on ripple discrimination. Here and below, lines represent classifier performance; solid: trials without light; dashed: trials with light. Inactivation of VIP (2-way ANOVA, f(1) = 7.45, p = 0.0072), PV (2-way ANOVA, f(1) = 27.50, p = 5.094e-6), and SST (2-way ANOVA, f(1) = 58.62, p = 1.98e-9) neurons all improved ripple discrimination performance. **F, G, H.** Effects of inactivation of VIP (F), PV (G), and SST (H) neurons on discrimination of deviant ripple X under REG versus RND contexts. VIP inactivation decreased classification performance (2-way ANOVA, f(1) = 10.058, p = 0.0019) while SST inactivation improved performance (2-way ANOVA, f(1) = 12.47, p = 0.001), and PV inactivation did not have a significant effect (2-way ANOVA, f(1) = 0.67, p = 0.42). **I, J, K.** Effects of inactivation of VIP (I), PV (J), and SST (K) neurons on discriminating background and deviant ripples under the REG context. All inactivations improved performance: PV (2-way ANOVA, f(1) = 50.18, p = 1.26e-8); SST (2-way ANOVA, f(1) = 58.45, p = 2.05e-9); VIP (2-way ANOVA, f(1) = 20.54, p = 1.26e-5) neuronal inactivation. **L, M, N.** Effects of inactivation of VIP (L), PV (M), and SST (N) neurons on discriminating background and deviant ripples in the RND context. Both PV (2-way ANOVA, f(1) = 29.35, p = 2.91e-6) and SST neuronal inactivation (2-way ANOVA, f(1) = 75.31, p = 7.93e-9) enhanced performance, while VIP neuronal inactivation had no effect on classification performance (2-way ANOVA, f(1) = 3.02, p = 0.085). **O, P, Q.** Difference between classification performance with and without light inactivation for corresponding panels I-N. VIP inactivation (2-way ANOVA, f(1) = 9.8941, p = 0.0020) and PV inactivation (2-way ANOVA, f(1) = 14.407, p = 0. 0048) had significant differences in their effects on SVM discrimination performance across REG and RND contexts while SST inactivation did not (2-way ANOVA, f(1) = 1.5055, p = 0.23).

First, to establish a baseline in ripple discriminability based on the tuning stimulus, we computed the effect of inactivation of inhibitory neurons on discriminability between two ripples (Figure 8A left). PV and SST inactivation led to an increase in ripple discriminability, reaching higher than 90% discriminability at n = 200 neuron population size (2-way ANOVA, f(1) = 27.50, p_PV_ = 5.094e-6, Figure 8D; 2-way ANOVA, f(1) = 58.62, p_SST_ = 1.98e-9, Figure 8E). VIP inactivation had a significant, but much weaker effect (2-way ANOVA, f(1) = 7.45, p = 0.0072, Figure 8C, D, E). These results are consistent with prior studies that found that PV and SST neurons affect neuronal tuning properties (Aizenberg et al., 2015; Li et al., 2015; Natan et al., 2015, 2017; Phillips and Hasenstaub, 2016; Kato et al., 2017; Phillips et al., 2017), whereas VIP neurons have an inconsistent effect on tuning in AC (Mesik et al., 2015; Tobin et al., 2024).

Next, we compared the SVM performance in discriminating the deviance response under REG versus RND contexts. If VIP neurons play a selective role in providing context-dependent information, we expected that their inactivation would impair the discriminability of deviance responses across REG and RND contexts. By contrast, for PV and SST neurons, we expected that their inactivation would have little effect on context discriminability. Indeed, we found that inactivating VIP neurons impaired context discriminability of deviance responses, which is consistent with a decrease in the difference in deviance responses under the two contexts (2-way ANOVA, f(1) = 10.058, p = 0.0019, Figure 8F). PV inactivation did not have a significant effect on context discriminability (Figure 8G), whereas SST inactivation improved the discriminability performance (2-way ANOVA, f(1) = 12.47, p = 0.001, Figure 8H).

Finally, we asked whether deviance detection under the two contexts improved with inhibitory neuronal inactivation. Surprisingly, we found that at a population level, inactivating all inhibitory neuron classes -- VIP, PV and SST neurons -- improved deviance detectability in REG context (2-way ANOVA, f(1) = 20.54, p_VIP_ =1.26e-5, 2-way ANOVA, f(1) = 50.18, p_PV_ = 1.26e-8, 2-way ANOVA, f(1)=58.45, p_SST_ =2.05e-9, Figure 8I, J, K). In RND context, VIP inactivation did not have an effect, whereas PV or SST inactivation also drove an improvement in deviance detection (2-way ANOVA, f(1) = 29.35, p_PV_ = 2.90e-6, 2-way ANOVA, f(1) = 75.31, p_SST_ = 7.92e-11, Figure 8 L, M, N). The effect PV or SST inactivation was strongest around a population of n = 10-25 neurons (Figure 8P, Q), suggesting that just a relatively small number of neurons can be involved in deviance detection. By contrast, the effect of VIP inactivation on deviance detection in REG context increased with increasing number of neurons (Figure 8O), suggesting that VIP neurons transmit the information about context to larger cortical populations. The finding that deviance detection at population level is increased when any of the three inhibitory neuron types are inactivated suggest that the cortical circuits operate in suboptimal regime in everyday hearing, potentially allowing for context-dependent modulation by different inhibitory neuronal classes (Tobin et al., 2024). This modulation is specific to regular context only for VIP neurons, in line in our analysis of their differential effect on single neuronal responses (Figure 6).

## DISCUSSION

In this study, we investigated how statistical regularity in sound sequences influences cortical responses to deviant auditory stimuli, and how inhibitory neurons contribute to this process. Using calcium imaging and optogenetic inactivation in awake mice, we found that auditory cortex (AC) neurons exhibited stronger deviance responses when deviant sounds followed a regular context compared to a random one. This enhancement was not uniform across cell types: while PV and SST interneuron inactivation broadly increased deviance responses with mixed effect of context, inactivation of VIP neurons selectively reduced deviance responses in the regular context. These findings reveal a distinct role for VIP neurons in modulating context-dependent deviance detection in AC.

Our results align with human studies showing that deviance detection is enhanced by temporal predictability in sound streams. Previous EEG and behavioral work demonstrated that listeners respond more robustly to violations in regular versus random sequences, but the underlying neural circuitry was unknown (Barascud et al., 2016; Sohoglu and Chait, 2016; Southwell and Chait, 2018). The stimuli used in these studies were comprised of pure tones presented in sequences with varied temporal statistics, either random or regular. While our stimuli use complex sounds (moving ripples) as the building blocks, the main stimulus manipulation we investigated is still changes to the temporal context. By probing neuronal subtypes in mouse AC, we provide direct evidence for circuit-level modulation of deviant responses by context. Our results suggest that temporal regularity enhances deviance detection not merely by boosting adaptation or prediction error in individual neurons (Ulanovsky et al., 2003, 2004; Natan et al., 2015, 2017; Parras et al., 2017), but through targeted engagement of specific inhibitory networks.

A key contribution of this work is identifying VIP neurons as selective modulators of context-dependent deviance responses. While VIP neurons have been implicated in disinhibitory circuits across sensory cortical regions (Mesik et al., 2015; Pakan et al., 2016; Kuchibhotla et al., 2017; Cone et al., 2019; Millman et al., 2020; Myers-Joseph et al., 2024; Tobin et al., 2024) and have been implicated in oddball and unexpected stimulus responses (Yarden et al., 2022; Bastos et al., 2023; Ferguson et al., 2023; Najafi et al., 2025), their role in encoding statistical structure in naturalistic auditory stimuli was previously untested. We show that VIP inactivation abolished the enhancement of deviance responses in the regular context without affecting responses in the random context. This selective effect suggests that VIP neurons integrate contextual information—potentially from top-down projections (Askew et al., 2019; Apicella and Marchionni, 2022; Ferguson et al., 2023) — to gate prediction error signaling (Friston and Kiebel, 2009; Friston, 2010; Heilbron and Chait, 2018). In contrast, PV and SST neurons modulate gain and adaptation more uniformly, regardless of context, consistent with their established roles in first-order sensory processing.

Our findings also have implications for how auditory scenes are represented at the population level. Using SVM-based decoding, we found that context information was more easily extracted from single-neuron responses in the regular condition, while population-level decoding performance plateaued more quickly. This shift toward a distributed, redundant code in regular contexts may reflect a network reorganization mediated by VIP neurons, as previously shown in auditory cortex (Tobin et al., 2024). Interestingly, inactivation of all interneuron types improved decoding of deviant stimulus identity, suggesting that inhibition refines but also constrains the dynamic range of population responses.

One strength of our approach is the use of spectrally rich ripple stimuli, which elicit reliable responses in a broad population of AC neurons. These stimuli better approximate the complexity of natural sounds than pure tones and may engage higher-order auditory computations relevant to speech, communication, and environmental navigation. Our observation that deviance detection operates robustly across both excitatory and inhibitory populations reinforces the idea that prediction-related computations are a core function of auditory cortex.

This work also opens new questions about the origins of context signals that modulate VIP activity. VIP neurons are known to receive input from neuro-modulatory and associative cortical areas, including prefrontal cortex (Kuchibhotla et al., 2017; Askew et al., 2019; Apicella and Marchionni, 2022; Ferguson et al., 2023; Ramamurthy et al., 2023). The selectivity of VIP modulation for regular contexts may arise from top-down prediction signals that are absent in random sequences. Future studies using pathway-specific manipulations or recordings in interconnected regions will be needed to determine how these inputs shape VIP function (Furutachi et al., 2024).

Several caveats warrant consideration. First, while we focused on layer 2/3 pyramidal neurons, different interneuron subtypes span cortical layers and exhibit layer-specific functions. Our widefield optogenetic inactivation likely affected VIP, PV, and SST neurons in multiple layers, and the net effect observed in superficial recordings may reflect combined contributions from distinct circuits (Meyer et al., 2011; Tremblay et al., 2016; Yavorska and Wehr, 2016; Naka et al., 2019; Wu et al., 2022; David et al., 2023). Second, the precise nature of the statistical computations—adaptation versus prediction error—encoded by each subtype remains to be disentangled. Simultaneous recordings across AC layers and subcortical regions could further clarify how prediction errors are computed and propagated.

In our recordings, we observed a substantial fraction of neurons with negative calcium responses to repeated sounds (Figures 2, 5). Although less frequently emphasized in the literature, such suppressive calcium signals in auditory cortex have been reported by multiple groups(Kato et al., 2015; Solyga and Keller, 2025; Tsukano et al., 2025). Notably, ACx neurons exhibited net negative calcium responses to auditory stimuli while showing positive responses to visual inputs, suggesting a modality-specific phenomenon rather than a technical artifact(Solyga and Keller, 2025). The mechanisms underlying these negative signals remain incompletely understood. One possibility is that calcium imaging captures inhibitory neurons more comprehensively than extracellular electrophysiology, which is biased toward larger excitatory neurons, though our prior labeling experiments indicate inhibitory cells are only a minority of the imaged population (Tobin et al., 2024). Furthermore, with habituation to sounds over days (which we perform in our experiments), the relative fraction of suppressive responses increases (Kato et al., 2015). Facilitating this may be a non-linear transfer function between spiking and GCaMP fluorescence, such that decreases in activity produce larger or more detectable fluorescence changes than modest increases in spiking. Regardless of their precise origin, our main conclusions are robust to response polarity: when analyses are restricted to neurons with positive responses, the key effects—particularly the context-dependent deviance responses—remain unchanged. These observations underscore the importance of reporting the full range of neuronal responses and highlight the need for future work combining intracellular recordings with calcium imaging to resolve the spiking-to-fluorescence relationship in suppressive responses.

In summary, our study demonstrates that statistical context modulates deviance detection in auditory cortex through specific inhibitory circuits. We identify inhibitory neurons, and specifically VIP neurons, as key players in shaping deviant responses based on contextual statistical structure. These findings support models of predictive coding in sensory cortex and suggest that context-sensitive disinhibition is a fundamental mechanism for flexible sensory processing. By revealing how different inhibitory neurons contribute to deviance detection under structured versus random auditory regimes, our results advance understanding of the neuronal basis of auditory perception in complex environments.

## Acknowledgements

The authors would like to thank the members of the Geffen laboratory and Yale Cohen for discussions and helpful comments. This work was supported by the following grants from NIH: R01DC15527, R01DC014479, R03DC013660, R01NS113241 to MNG, and K99DC019504 to KCW. XD was supported by T32DC016903.

