## Supplementary Figures for "The role of inhibitory neurons in deviance sound detection in regular and random statistical contexts"

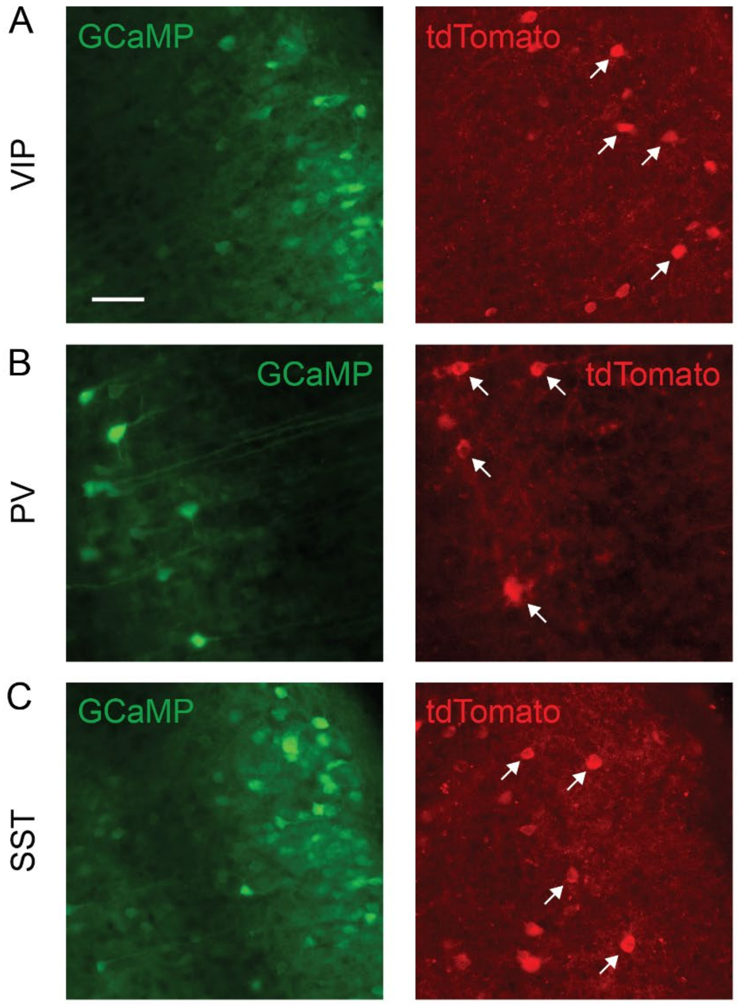


**Supplementary Figure S1. Expression of tdTomato in auditory cortex.**
Example immunohistochemistry images of GCaMP (left) and tdTomato (right) expression in auditory cortex. TdTomato expression was amplified using the Rockland RFP primary antibody. **A.** Example image from a VIP-cre mouse, with GCaMP on the left and tdTomato on the right. Arrows indicate example cells labeled with tdTomato. **B.** Similar example from a PV-cre mouse. **C.** Similar example from a SST-cre mouse.

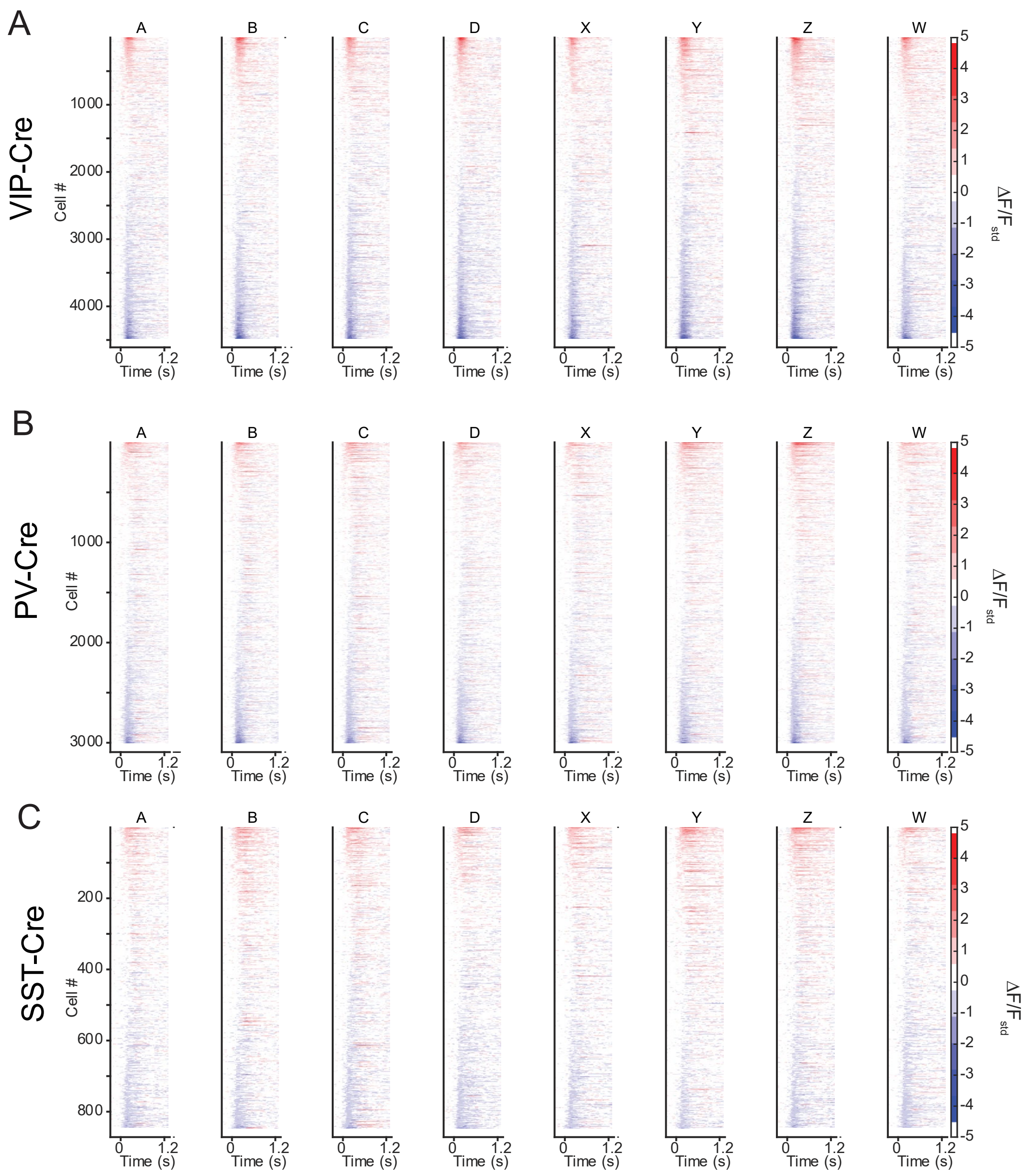


**Supplementary Figure S2. Distribution of the mean responses to each of the 8 ripples** when presented as a tuning stimulus across the population of recorded neurons for VIP-Cre (**A**), PV-Cre (**B**), and SST-Cre (**C**) mice. Color depicts the relative fluorescence.

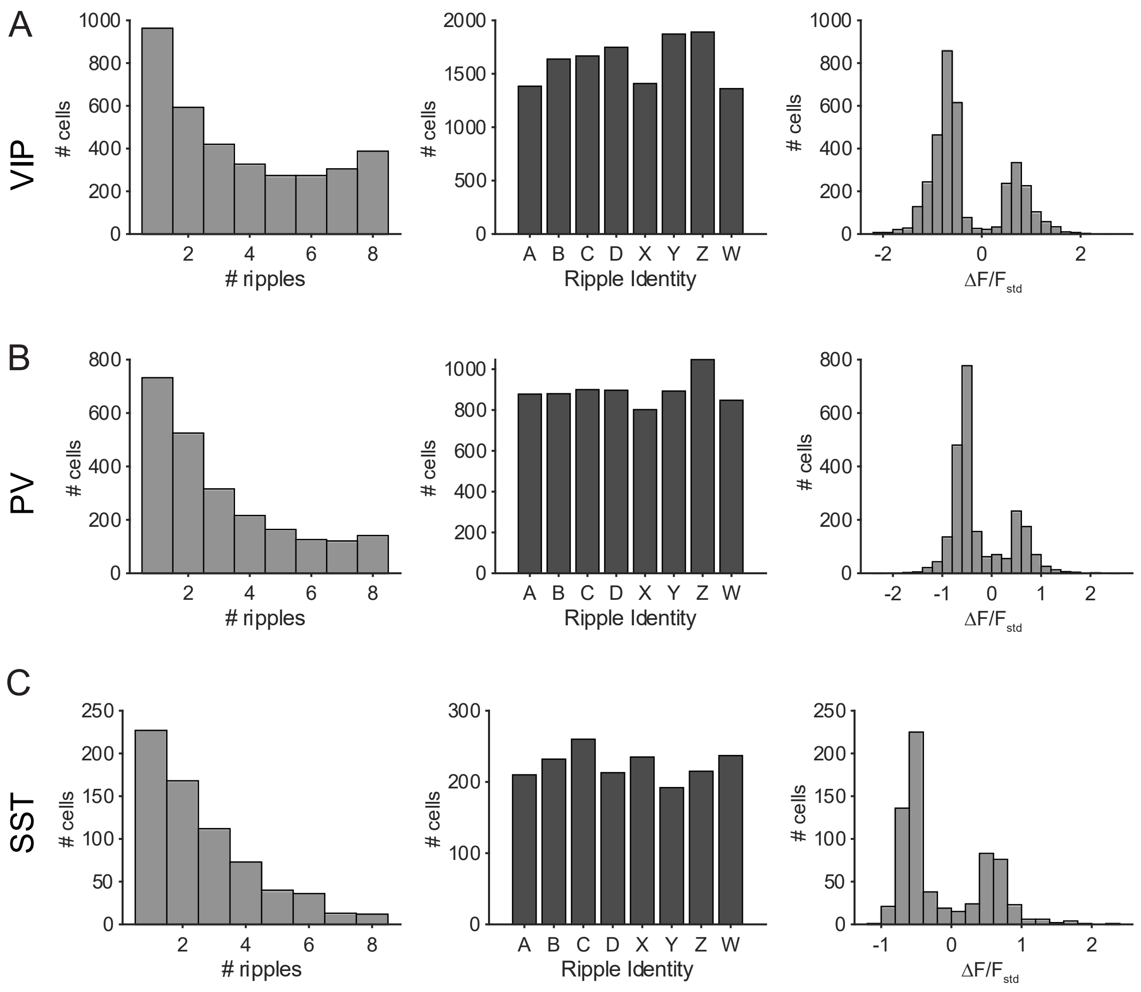


**Supplementary Figure S3. Distribution of responses to ripple sounds across recorded neuronal populations** for VIP-Cre (**A**), PV-Cre (**B**), and SST-Cre (**C**) mice. Left: Histogram of the number of neurons that are significantly responsive to different number of ripples. Note that the neurons exhibited specificity in responses with most neurons responding to at most 1 ripple. Center. Histogram of the number of neurons significantly responsive to each of the 8 ripples. Right. Histogram of response strength across different ripples. We did not observe systematic differences in responsiveness to different ripples across the cell lines.

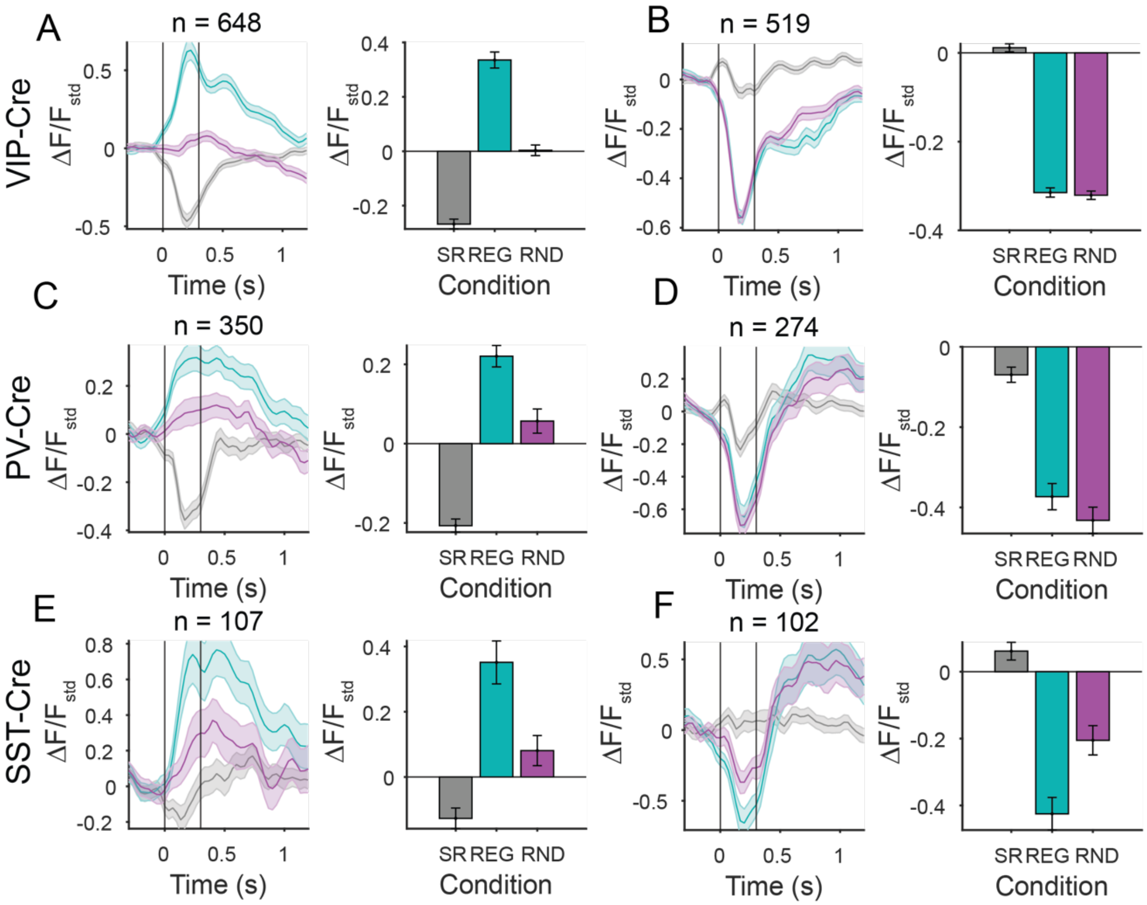


**Supplementary Figure S4. Consistent effects for positive, but not negative differential novelty responses across mouse lines. A-F**. Left: Time course of responses to the deviant ripple in regular (cyan), random (magenta) or baseline (gray) contexts across neurons. Right: Mean responses to the deviant ripple in regular, random and baseline contexts. Error bars: ste. **A**. VIP-Cre mice, n = 648 neurons with significantly context-dependent positive deviance responses (linear mixed effects model, $F(1,1294)=103.96$, $p = 1.56e-23$). **B**. VIP-Cre mice, n = 519 neurons with significantly context-dependent negative deviance responses (linear mixed effects model, $F(1,1036)=0.98$, $p = 0.32$). **C**. PV-Cre mice, n = 350 neurons with significantly context-dependent positive deviance responses (linear mixed effects model, $F(1,698)=16.26$, $p = 6.12e-5$). **D**. PV-Cre mice, n = 274 neurons with significantly context-dependent negative deviance responses (linear mixed effects model, $F(1,546)=1.95$, $p = 0.16$). **E**. SST-Cre mice, n = 107 neurons with significantly context-dependent positive deviance responses (linear mixed effects model, $F(1,212)=11.40$, $p = 0.0009$). **F**. SST-Cre mice, n = 102 neurons with significantly context-dependent negative deviance responses (linear mixed effects model, $F(1,202)=11.38$, $p = 0.00089$).

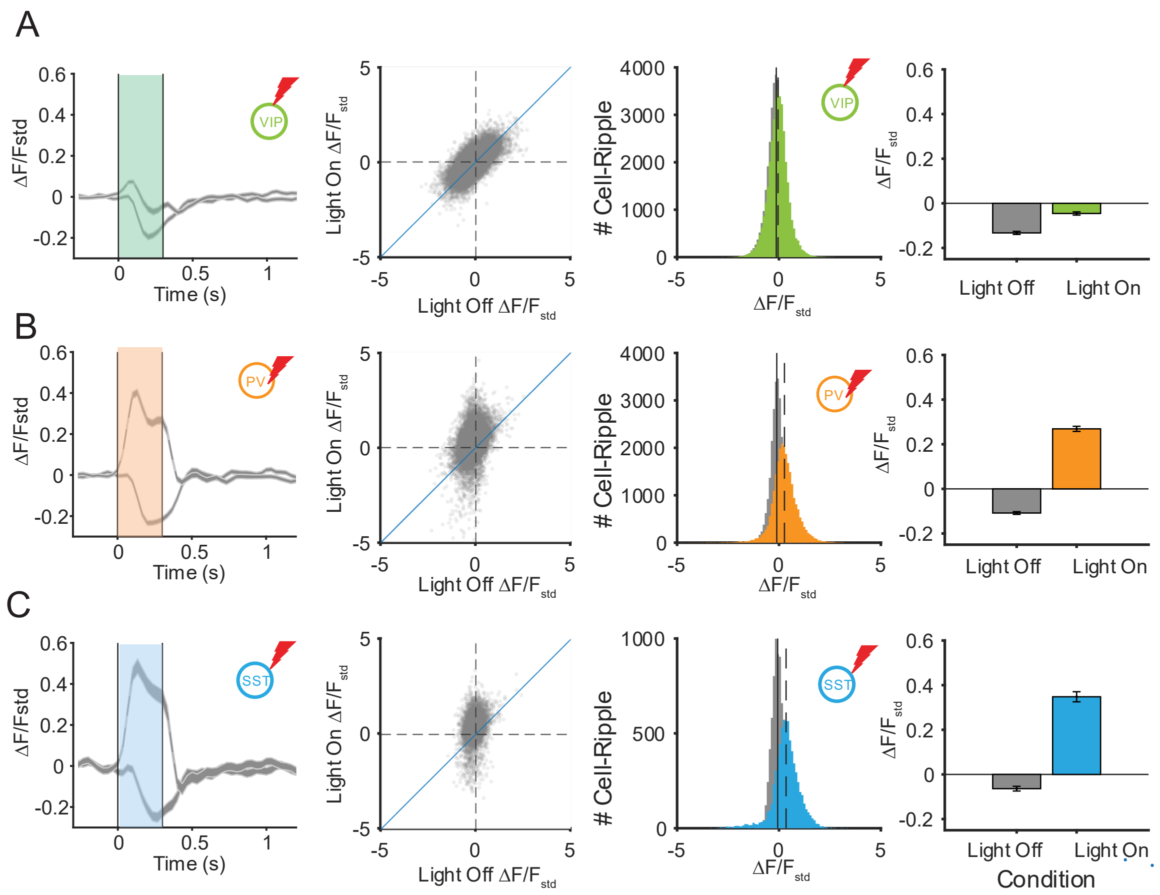


**Supplementary Figure S5. Suppressing neuronal activity only partially affected the responses of cortical neurons to dynamic moving ripples. A**. Effects of VIP inactivation. Left: Timecourse of responses to ripple X across the population of neurons on light on (dashed gray) and light on (solid gray) trials. Center left: Mean response of neurons to each of 8 ripples on light on versus light off trials. Each dot depicts a neuron-ripple combination. Center right. Histogram of responses to each of 8 ripples on light on (colored) and light off (gray) trials. Right: Mean response to ripples on light off (gray) and light on (colored) trials. N = 4609, p = $5.87*10^{-135}$. **B**. Effects of PV inactivation. Axes same as in **A**. N = 4093, $p < 1*10^{-20}$. **C**. Effects of SST inactivation. N = 871, $p <1*10^{-20}$. Axes same as in **B**.

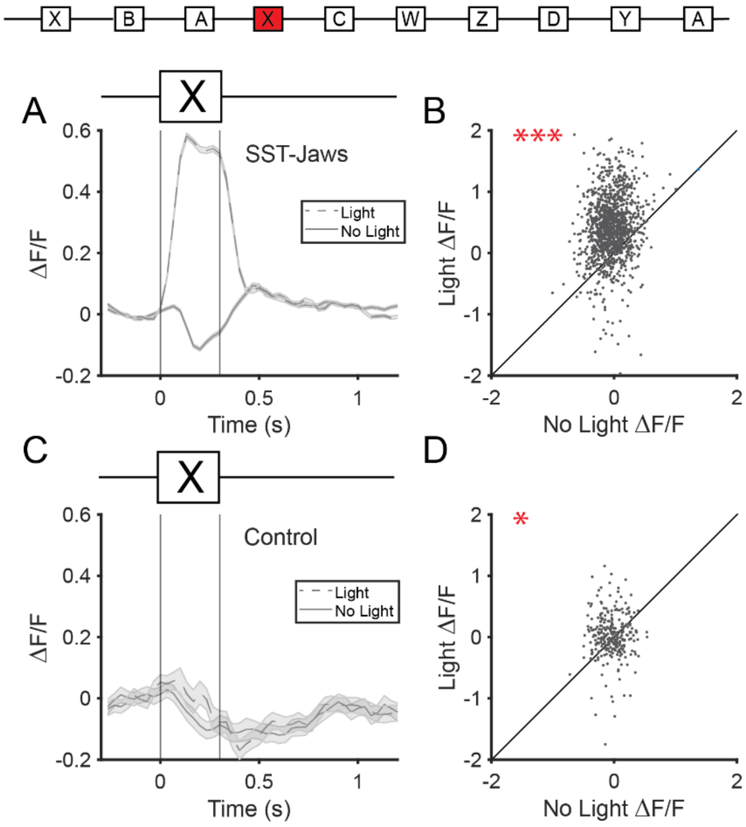


**Supplementary Figure S6. Effects of light activation in controls.**

**A.** Sound response of all neurons injected with the JAWS-tdTomato virus cocktail with (dashed line) and without (solid line) optogenetic manipulation. **B.** Scatter plot of the mean responses of the first 300 ms post-stimulus onset for each neuron in A. There is a significant effect of the light (mean = -0.455, linear mixed effects model, $F(1, 2978) = 794.45$, $p = 3.88e-155$). **C, D.** Similar plots as in A,B but for neurons from mice only injected with tdTomato and no JAWS. There no is significant effect of light activation (mean = -0.06, linear mixed effect model, $F(1, 2978) = 3.24$, $p = 0.72$). Furthermore, there is a significant difference in effect between the light effect in responses recorded from neurons injected with JAWS and control virus (mean = -0.39, linear mixed effect model, $F(1, 2978) = 114.098$, $p = 3.67e-26$).

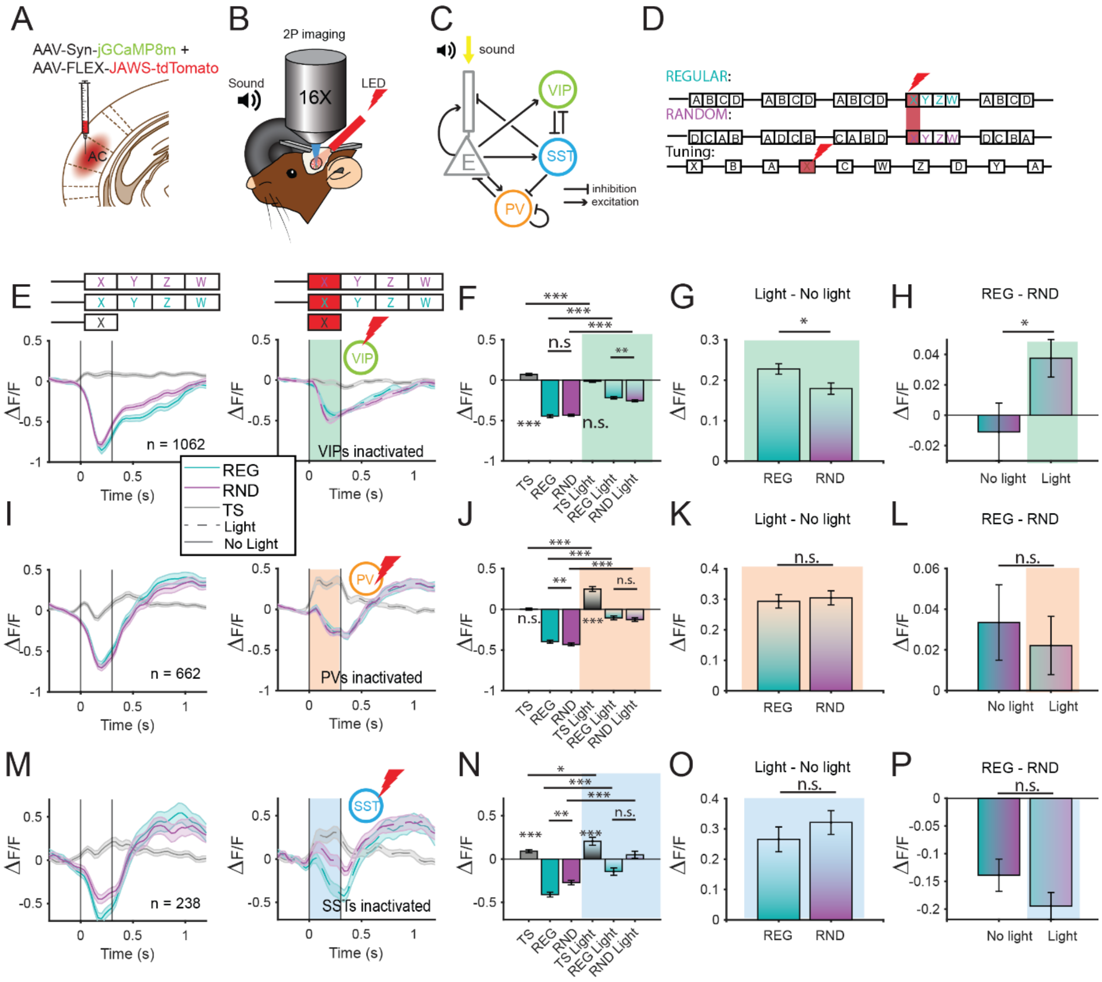


**Supplementary Figure S7. Neurons with a negative novelty response exhibit differential effects under optogenetic inactivation of three types of inhibitory neuron classes.**

**A-P.** Same format as Figure 6. **E, F, G, H.** Effects of optogenetic inactivation of VIP neurons on neurons with a negative novelty response. VIP inactivation increases novelty responses (linear mixed effect model, $F(1, 4244) = 196.62$, $p = 1.06e\text{-}43$) while not significantly differential effect modulating responses based on context (linear mixed effects model, $F(1, 4244) = 2.78$, $p = 0.095$). Neurons responsive to VIP inactivation exhibited a significant baseline sound response without light (t-test, $p = 1.9e\text{-}07$) but not with (t-test, $p = 0.27$). **I, J, K, L.** Inactivation of PV neurons increased negative responses (linear mixed effects model, $F(1, 2644) = 112.92$, $p = 7.42e\text{-}26$), but exhibited no differential effect of context (linear mixed effects model, $F(1, 2644) = 1.36$, $p = 0.244$). Neurons responsive to PV inactivation did exhibit a significant baseline sound response with (t-test, $p = 7.42e\text{-}17$) but not without (t-test, $p = 0.87$) light. **M, N, O, P.** Inactivation of SST neurons increased novelty responses (linear mixed effects model, $F(1, 948) = 74.88$, $p = 2.12e\text{-}17$), but not differentially based on context (linear mixed effects model, $F(1, 948) = 0.67$, $p = 0.41$). Neurons responsive to SST inactivation did exhibit a significant baseline sound response with (t-test, $p = 1.22e\text{-}05$) and without (t-test, $p = 8.60e\text{-}07$) light.
